# Phylogenomics provides insights into the evolution of cactophily and host plant shifts in *Drosophila*

**DOI:** 10.1101/2022.04.29.490106

**Authors:** Nicolás Nahuel Moreyra, Francisca Cunha Almeida, Carson Allan, Nicolás Frankel, Luciano Matías Matzkin, Esteban Hasson

## Abstract

Cactophilic species of the *Drosophila buzzatii* cluster (*repleta* group) comprise an excellent model group to investigate genomic changes underlying adaptation to extreme climate conditions and host plants. In particular, these species offer a subject to study the transition from chemically simpler breeding sites (like prickly pears of the genus *Opuntia*) to chemically more complex hosts (columnar cacti). Here, we report four highly contiguous genome assemblies of three species of the *buzzatii* cluster. Based on this genomic data and inferred phylogenetic relationships, we identified candidate taxonomically restricted genes (TRGs) likely involved in the evolution of cactophily and cactus host specialization in internal branches of the subgenus *Drosophila*. Functional enrichment analyses of TRGs within the *buzzatii* cluster identified genes involved in detoxification, water preservation, immune system response, anatomical structure development, and morphogenesis. In contrast, processes that regulate responses to stress, as well as the metabolism of nitrogen compounds, transport, and secretion were found in the set of species that are columnar cacti dwellers. These findings are in line with the hypothesis that those genomic innovations brought about instrumental mechanisms underlying adaptation in a group of species that speciated in the arid regions of South America.

## 1. INTRODUCTION

Comparative genomics provides invaluable information for the study of organismal biology, the evolution of genes and gene families, and phylogenetic relationships (Koonin et al., 2000; Hardison, 2003; Miller et al., 2004; Gabaldón, 2008). Fortunately, genome sequencing technologies are producing genomic data from thousands of non-model organisms (i5K Consortium, 2013; Thomas et al., 2020; Kim et al., 2021) leading to new hypotheses about major evolutionary events (Almudi et al., 2020). Insect diversification is an event that has fascinated evolutionary biologists. It is intrinsically related to the conquest of new environments (Grimaldi et al., 2005), and provides a great subject to study genomic changes driving adaptation and the evolution of innovations that facilitate ecological transitions.

One interesting outcome of comparative genomic studies is the discovery of substantial variation in gene number across genomes of related species, denoting the existence of species-specific genes (Clark et al., 2007; Hou & Lin, 2009). In this context, the concept of gene homology is central when comparing genomes. Homologous genes are defined as those that derive from a common ancestor, and sequence similarity is the standard criterion for establishing homology (Kuzniar et al., 2008). Sequence similarity, however, may be considered as a working hypothesis for homology that should be confirmed by further evidence such as conserved synteny (Webber & Ponting, 2004; Vakirlis et al., 2020). Proper homolog identification in whole genome studies is not only necessary for phylogenetic reconstruction, but also of utmost relevance to understanding patterns of gene content and functional conservation throughout the tree of life (Almudi et al., 2020; Fernández & Gabaldón, 2020). Genes lacking detectable homologs anywhere else in the tree of life are known as orphans and are quite frequent in almost any organism (Wilson et al., 2005; Khalturin et al., 2009; Singh & Wurtele, 2020).

How orphan genes originate, what their roles are, and what forces drive their evolution are open questions (Tautz & Domazet-Lošo, 2011; Singh & Wurtele, 2020). Evidence suggests that orphan genes evolve quickly after emerging either by sequence divergence from preexisting genes or *de novo* (gene birth) (Tautz & Domazet-Lošo, 2011; Tautz, 2014; Singh & Wurtele, 2020; Vakirlis et al., 2020), although they can also arise from gene losses in external branches of the phylogeny or from horizontal gene transfer (Dunning Hotopp, 2011; Tautz & Domazet-Lošo, 2011). Moreover, the concept of orphan can be extended to genes that can only be found in a small group of related species or lineage-specific genes, which are known as taxonomically restricted genes (TRGs) (Wilson et al., 2005; Khalturin et al., 2009). The evolution of TRGs has been proposed to be the underlying genetic basis of adaptive evolutionary innovations, with functions involved in the interaction with the environment, and, thus, as part of mechanisms involved in the acquisition of new niches (reviewed in Johnson, 2018).

The family Drosophilidae has been widely studied because of remarkable features that facilitate the study of ecology, development, taxonomy, divergence, and phylogenetic relationships (Kim et al., 2021). The ecology of *Drosophila* is highly diverse, including species that breed on fruits, cacti, flowers, mushrooms, and tree saps (Markow & O’Grady, 2008). Likewise, the wide distribution of this genus offers a range of phenotypes associated with its diverse ecology. For instance, host shifts from fruits to cacti as well as the adaptation to arid and desertic environments have been instrumental in the evolution of the *repleta* group (Markow & O’Grady, 2008; Oliveira et al., 2012). The ability to utilize necrotic cacti as breeding substrates, is an evolutionary novelty that arose independently at least twice and allowed the spread and diversification of the *repleta* group in American arid lands (Oliveira et al., 2012). Cactophilic species of the *repleta* group can be grossly divided into two groups on the basis of the type of host plant use: species that breed on prickly pears (genus *Opuntia*, subfamily Opuntioideae) and columnar cactus (subfamily Cactotideae) breeders. The current evidence suggests that prickly pears, a generally more benign substrate for flies, are the ancestral state of host plant use in the *repleta* group (Oliveira et al., 2012; Hasson et al., 2019).

The *Drosophila*-cactus-yeast system has long been a model for ecological genetic studies (Barker & Starmer, 1982; Heed & Mangan, 1986; Barker et al., 2013; Fogleman & Danielson, 2001) involving, among others, the *D. mulleri* subgroup, which includes species distributed throughout the Americas. The North American species that are desert inhabitants comprise the *mulleri* complex (Oliveira et al., 2012) and represent a case study of adaptation to breeding on chemically hostile host plants (Heed & Mangan, 1986; Fogleman & Danielson, 2001). The *D. buzzatii* complex, the sister group of the *mulleri* complex, includes the *buzzatii, martensis*, and *stalkeri* clusters that evolved in South America and Caribbean Islands (Ruiz & Wasserman, 1993). The former is an ensemble of seven species: *D. antonietae, D. borborema, D. buzzatii, D. gouveai, D. koepferae, D. serido*, and *D. seriema*;all endemic to South America (Manfrin & Sene, 2006), except for *D. buzzatii* that reached a semicosmopolitan distribution in historical recent times (Fontdevila, 1989). Though morphologically very similar, species of the *buzzatii* cluster can be distinguished by male genital morphology (Manfrin & Sene, 2006) and fixed and polymorphic chromosomal inversions (Ruiz et al., 2000). Based on these criteria, the cluster has been divided into two groups, one including *D. buzzatii* and the the *serido* sibling set comprising the remaining species (Manfrin & Sene, 2006). This classification has been corroborated using transcriptomic data (Hurtado et al., 2019), but has also been challenged by a mitogenomic study pointing to a different evolutionary history (Moreyra et al., 2019). In any case, relationships within the *serido* sibling set remain unclear.

Concerning patterns of host plant use in the *buzzatii* cluster, *D. buzzatii* has been mainly recovered from necrotic cladodes of several *Opuntia* species and marginally from columnar cacti, whereas the opposite has been observed in *D. koepferae* (Hasson et al., 2019). The other members of the cluster are mainly associated with columnar cacti (Manfrin & Sene, 2006). Experimental host shifts from chemically benign opuntias to the hostile environment of alkaloid-rich columnar cacti result in a decrease in survival, lengthening of development and increase in developmental instability in the *Opuntia* feeder *D. buzzatii*, whereas the columnar dweller *D. koepferae* fares better in cardón (columnar cacti) than in prickly pears (Hasson et al., 2019). Moreover, it has been shown that changes in gene expression are orchestrated in response to different factors: cactus alkaloids in *D. buzzatii* and alternative host plants in *D. koepferae* (Hasson et al., 2019; De Panis et al. submitted). These findings along with the fact that the *buzzatii* cluster comprises species at different stages of divergence, make it an excellent model to study the adaptive mechanisms underlying cactophily and host plant use specialization (reviewed in Hasson et al., 2009, 2019). However, most species of the *repleta* group that have been sequenced so far are from North America (Clark et al., 2007; Sanchez-Flores et al., 2016; Rane et al., 2019; Jaworski et al., 2020; Kim et al., 2021). Thus, to understand the genomic changes associated with the acquisition of cactophlily and host shifts from chemically simpler hosts like prickly pears to chemically complex columnar cacti, new genomic data are necessary, particularly from South American species.

Here we present the assembly and annotation of four new genomes of three cactophilic species of the *buzzatii* cluster and the re-annotation of the genome of *D. buzzatii*. Using genomic data of nine related *Drosophila* species we report the most complete phylogeny for the *buzzatii* cluster to date, divergence time estimates at each node, and candidate TRGs in all branches of the tree. We also focus on the molecular evolution of candidate TRGs that might be associated with cactophily in the *repleta* group and adaptation to the chemically stressful columnar cacti within the *buzzatii* cluster.

## 2. MATERIALS AND METHODS

### 2.1 Species selection

We sequenced four genomes of three species of the *buzzatii* cluster. Adult flies of single inbred lines of *D. antonietae* (strain MG.2, Argentina) and *D. borborema* (BOR, Brazil, Stock Center; #(BGS) 3403.4), and two lines representative of allopatric populations of *D. koepferae* from Argentina and Bolivia, strains Ko7.1 (DkoeA) and Ko11 (DkoeB), respectively, were selected for whole genome sequencing. Genomic data of seven other members of the *repleta* group retrieved from public databases were included in our study: *D. buzzatii, D. arizonae, D. mojavensis*, and *D. navojoa* (*mojavensis* cluster), and *D. aldrichi* as representatives of the *mulleri* complex (*mulleri* subgroup); and *D. hydei* and *D. mercatorum* of the *hydei* and *mercatorum* subgroups, respectively. Within the *repleta* group, species that belong to the *mulleri* subgroup are cactophilic whereas *D. hydei* and *D. mercatorum* are dietary generalists with the ability of feeding upon rotting fruits, vegetables, and cacti. Note that the *D. mercatorum* genome included in our study was originally reported as *D. repleta* by Rane et al. (2019) and assigned to the proper species by (Li et al., 2021). The genomes of *D. virilis* (*virilis* group, subgenus *Drosophila*), sister of the *repleta* group, and *D. melanogaster* as the only representative of the subgenus *Sophophora* were also included in the study (Throckmorton, 1975; Clark et al., 2007). Full information about genome assembly accession numbers and versions used for each species is presented in Table S1, and taxonomical and systematics information for each species is shown in Text S1.

### 2.2 Sequencing protocol

Genomes were sequenced following a hybrid approach that involved short and long reads technologies. First, Illumina Hiseq 2000 was employed to sequence paired-end reads at Centre Nacional d’Analisi Genomica (Barcelona, Spain; https://www.cnag.eu/) and at Centre de Regulació Genòmica (Barcelona, Spain; https://www.crg.eu/). Second, Pacific Biosciences (hereafter PacBio) long reads were sequenced in two stages. Initially, we sequenced the genomes of *D. borborema* and *D. koepferae* A in two SMRT P6/C4 cells using RS II technology at DNA Sequencing Core (University of Michigan, Michigan, USA; https://www.seqcore.brcf.med.umich.edu). Next, the genomes of *D. koepferae* A and B, *D. borborema*, and *D. antonietae* were sequenced using one SMRT cell for each with Sequel I technology at Arizona Genomics Institute (School of Plant Sciences, University of Arizona, Arizona, USA; https://www.genome.arizona.edu). Protocols of DNA extraction can be found in Text S2.

### 2.3 Quality control and filtering of reads

Quality of Illumina paired-end reads was analyzed with FastQC ver. 0.11.3 (Andrews, 2010). We only kept reads that had a mean Phred score (Q) > 25. Cutadapt ver. 1.16 (Martin, 2013) was applied to detect and extract remnant adapters from reads, and only those longer than 25 bp were retained. Then, Trimmomatic v0.33 (Bolger et al., 2014) was applied to remove reads with mean Q ≤ 25 using a sliding window approach. Reads with at least 20 bp were retained. PacBio long reads were analyzed to calculate the length distribution but no filter was applied given the base correction and polishing steps employed in the assembly protocol (see below).

### 2.4 Genome assembly

To assemble the four genomes we followed a *de novo* hybrid approach adapted from (Jaworski et al., 2020). First, a low heterozygosity genome assembly was obtained with Platanus (Kajitani et al., 2014) using Illumina paired-end reads. Second, DBL2OLC (Ye et al., 2016) was used to generate another assembly based on both Illumina and PacBio reads and the high confidence sequences (contigs and scaffolds) previously assembled with Platanus. PacBio reads were also used as input in a third genome assembly using Canu Assembler ver. 1.7 (Ye et al., 2016; Koren et al., 2017). This assembly consisted of correction, trimming, and assembly stages using almost all parameters by default (*correctedErrorRate* was set to 0.075). Third, a polishing method was individually applied to the assemblies generated with DBG2OLC and Canu. To achieve this, Illumina reads were mapped onto each genome assembly using Pilon ver. 1.22 (Zelle et al., 2014) to correct remaining sequencing errors, and the Arrow consensus caller (SMRT link ver. 3.0.2, (https://github.com/PacificBiosciences/GenomicConsensus) was utilized to detect and remove miss-assemblies by mapping the PacBio reads. After polishing, both assemblies were combined using Quickmerge ver. 0.2 (Chakraborty et al., 2016). For this purpose, the assembly obtained with DBG2OLC was employed as a reference in the genome alignment and the anchor length was set to the N50 value of the assembly built using Canu. Lastly, another polishing round was carried out on the merged assembly. The complete scheme of this protocol can be found in Figure S1.

Assembly contiguity was assessed with Quast ver. 4.6.3 (Gurevich et al., 2013) and completeness was evaluated using BUSCO ver. 4.1.4 (Seppey et al., 2019) for 3285 dipteran universal single-copy orthologs (BUSCO groups) obtained from OrthoDB ver. 10.1 (Kriventseva et al., 2019).

### 2.5 Genome annotation

To mask genomes before gene annotation, repetitive element identification and classification were performed following the advanced repeat library construction tutorial of MAKER ver. 2.31.10 (Holt & Yandell, 2011) (the full protocol description can be found in Text S3). Briefly, we *de novo* identified miniature inverted transposable elements (MITEs) as well as recent and old divergent long terminal repeats (LTRs) on each genome. We then masked each genome with its specific repeat library to search for new repeat elements that were missannotated before, using RepeatModeler ver. 1.0.11 (https://github.com/Dfam-consortium/RepeatModeler) with default parameters. The new unknown elements were searched against the transposase database using BLASTX ver. 2.9.0+ (Camacho et al., 2009), and reclassified as ‘known’ if significant matches (*e-value* < 1×10^-10^) to a transposon superfamily were found. All repeat sequences collected at this stage were compared to a *Drosophila* protein database downloaded from FlyBase release FB2019_01 (Thurmond et al., 2019). Elements with significant hits to genes were removed with ProtExcluder ver. 1.2 (Campbell et al., 2014; Thurmond et al., 2019). After excluding all gene fragments, we generated combined species-specific libraries by combining MITEs, LTRs, and identified repeat elements.

Genome annotation was accomplished in four steps with MAKER ver. 2.31.10 (Holt & Yandell, 2011), which construct gene models for the longest transcript per gene. In the first step, each genome assembly was masked with the species-specific repeat library and, then, gene models were built based on the mapping of transcripts and protein evidence. The transcriptomes of *D. antonietae, D. borborema, and D. koepferae* A were used as transcript evidence (Hurtado et al., 2019). The transcriptomes of each species plus another of a related species were mapped by setting the parameters *est* and *altest*, respectively. For *D. koepferae* B, the same transcriptome of *D. koepferae* A was used as self transcript evidence. We also re-annotated the genome of *D. buzzatii* using the same methodology but masking the assembly with the species-specific repeat library reported in Rius et al. (2016) and using the first annotation reported in Guillén et al. (2014) as reference in the mapping of species-specific transcriptomes (Hurtado et al., 2019; Mensch et al., unpublished results). The protein evidence mapped to each genome involved a set of non-redundant known proteins that was created by combining the FlyBase protein database release FB2019_01 and the UniProtKB/Swiss-Prot database release 2018_11.

In the second step, SNAP ver. 2006-07-28 (Korf, 2004) and Augustus ver. 3.2.3 (Stanke et al., 2008) were trained to detect exons, splice sites, and UTR regions of each gene. We first trained SNAP to build gene models with an annotation edit distance (AED) value ≤ 0.25 (Eilbeck et al., 2009) and with a protein product of at least 50 amino acids long. We collected the resulting trained sequences with the 1000 bp flanking regions using *fathom*, and *forge* was subsequently employed to calculate training parameters. The hmm-assembler script was then applied with both training sequences and parameters to build the final gene models. Second, we extracted mRNA sequences from the gene models generated using protein and transcript evidence (first step) to train Augustus. We applied BUSCO ver. 3.0.2 (Simão et al., 2015) to re-annotate the extracted mRNA sequences using 2799 dipteran BUSCO groups obtained from OrthoDB ver. 9.1 (Zdobnov et al., 2017). This step aimed to generate species-specific models for these conserved genes as well as to give an idea of the completeness of the annotation. Thus, the BUSCO run was set to reannotate these genes using BLAST searches and the built-in HMM model of *D. melanogaster*. The initial gene models constructed were then used to train Augustus and, consequently, to produce species-specific HMM models that were employed in the last step in MAKER.

In the third step, a new round of MAKER annotation was run applying the evidencebased gene models, and both gene models and species-specific parameters predicted by SNAP and Augustus. Then, steps 2 and 3 were repeated iteratively in additional annotation rounds to improve predictive power. The number of rounds required for each species was determined by measuring gene annotation performance, i.e, number of gene models, mean gene length, AED distribution, and completeness of BUSCO groups. Then, the annotated protein sequences were compared to the UniProtKB and eggNOG (Huerta-Cepas et al., 2019) databases to remove genes encoding non-eukaryotic protein from annotation (see Text S4 for details). Finally, to evaluate annotation quality, we also calculated the distribution of the AED across all gene models.

### 2.6 Phylogenomic analyses

#### 2.6.1 Protein datasets construction

We created protein sets for 13 genomes by selecting only the longest protein sequence product per gene. However, given the lack of protein sequence products (protein fasta files) available for *D. aldrichi* and *D. mercatorum* annotations, we applied Transdecoder ver. 5.5.0 (https://github.com/TransDecoder/TransDecoder/wiki) to generate the corresponding protein sets. Open reading frames (ORFs) of at least 100 amino acids long were predicted with the *TransDecoder.Predict* algorithm and, to maximize sensitivity, only ORFs with homology to known proteins or to common protein domains were retained in the final set of proteins. The UniprotKB/Swiss-Prot release 2020_06 (BLASTP search, *e-value* =*1×10^-5^*) and Pfam-A release 33.1 (Hmmscan search) databases were used for homology searches. Subsequently, several Python *ad hoc* scripts were employed to create the subset with the longest protein product per gene for *D*. *aldrichi, D. arizonae, D. hydei, D. melanogaster, D. mojavensis, D. navojoa*, and *D. virilis*.

#### 2.6.2 Species phylogeny

Phylogenetic relationships were inferred using a set of dipteran BUSCO groups. Thus, the corresponding protein sequences for the 13 genomes included in our analyses were aligned (auto mode) using MAFFT ver. 7.215 (Katoh & Standley, 2013). Next, trimAl ver. 1.4.rev22 (Capella-Gutiérrez et al., 2009) was applied to remove poorly aligned regions in each case and, then, an amino acid sequence supermatrix was built by concatenating the alignments of each BUSCO group. The species tree was computed using IQ-TREE ver. 2.0.3 (Minh et al., 2020) in a maximum likelihood search with 1000 bootstrap replicates and automatically determining the best-fit substitution model for each partition (BUSCO group). Gene- and site-concordance factors (gCF & sCF) were calculated to investigate potential discordance across loci and sites (Minh et al., 2020). IQ-TREE was run again to estimate all single-locus trees and to calculate gCF and sCF values for each branch of the species tree. Bootstrap, gCF, and sCF values were then summarized and plotted onto the species tree using FigTree ver. 1.4.4 (Rambaut 2007).

#### 2.6.3 Divergence times

Divergence times were estimated by means of two approaches employing proteincoding sequence (CDS) alignments of BUSCO groups. In the first, we estimated divergence times using the neutral mutation rate empirically obtained for *D. melanogaster* (Keightley et al. 2009) to set up a strict clock. Only alignments with codon usage bias (CUB) lower than 0.375 (as in Obbard et al., 2012) were retained and, then, 4FDS were extracted and concatenated into the matrix. PartitionFinder2 (Lanfear et al., 2017) was run to estimate the substitution model that best fitted each BUSCO group alignment. Divergence times were estimated using BEAUti and BEAST ver. 1.10.4 (Drummond & Rambaut, 2007). BEAUti was first used to import the matrix and to specify the evolutionary substitution model for each partition. We also set as priors a Birth-Death process for speciation and a strict molecular clock with a molecular substitution rate of 3.46×10^-9^ (stdev = 0.281). Bayesian Inference searches were then run with BEAST by setting a MCMC run of 15 million generations with parameters logged every 1000 generations. Convergence of the chain was evaluated with Tracer ver. 1.7.1 (Rambaut et al., 2018) by discarding 10% of trees as burn-in. TreeAnnotator ver. 1.10.4 (available as part of the BEAST package) was applied to summarize the information of the recovered trees, and the annotated tree was visualized using FigTree.

In the second approach, we followed the procedure outlined in Suvorov et al. (2021) to generate a node age-calibrated phylogeny using MCMCTREE software, which is part of the PAML package ver. 4.9 package (Yang 1997, 2007). Firstly, the complete sequence alignments of BUSCO groups were concatenated without previous filters, and the resulting matrix was subsequently divided into 3 partitions corresponding to each codon position. Secondly, as MCMCTREE requires at least two time constraints and due to the lack of fossils or geological events to calibrate the ingroup of our study, we employed the estimates for the separation between the *Drosophila* and *Sophophora* subgenera (47 Mya with lower and upper bounds of 43 and 50 Mya) and for the *virilis-repleta* radiation (23-30 Mya) reported in Suvorov et al. (2021) as node age constraints. For this step, a GTR+G substitution model and a Birth-Death process for speciation were also applied and the remaining parameters were set as default. MCMCTREE was run to obtain maximum likelihood estimates of branch length, gradient, and Hessian matrix (which constructs an approximation to the likelihood function by Taylor expansion) (dos Reis & Yang, 2011) to approximate the likelihood for the three partitions. Then, to estimate divergence times, MCMCTree was run again over 60 million generations sampling parameters every 3000 generations and discarding 10% of the states as burn-in. Lastly, in order to check for convergence, we repeated the run and compared the results in Tracer. The time-calibrated tree was visualized using FigTree.

### 2.7 Ortholog gene evolution

#### 2.7.1 Ortholog inference

The identification of potential orthologs (orthogroups) across the 13 proteomes was conducted with OrthoMCL ver. 2.0.9 (Li et al., 2003), and the OrthoMCL Pipeline (https://github.com/apetkau/orthomcl-pipeline) was applied to automate this task. By using the sets of one protein per gene as input, BLASTP intra- and interspecific comparisons were done by setting the *e-value* to 1×10^-5^ and match cutoff to 50%. Identified orthogroups were analyzed to determine candidate lineage-specific orthologs, i.e. orphans (genes restricted to only one species) and TRGs (genes exclusive to one taxonomic group or clade), in the species phylogeny. For this purpose, we evaluated the presence of these kinds of genes at selected branches in the tree according to taxonomic classification, dietary preference, and primary host plant use. To this end, we evaluated: 1) the root of the tree including all species; 2) the subgenus *Drosophila*; 3) the *repleta* group; 4) the *mulleri* subgroup, all species that use cacti as breeding and feeding resources; 5) the *mulleri* complex, encompassing the four North American cactophiles, i.e. *D. aldrichi* plus the species of the *mojavensis* cluster; 5) the *buzzatii* complex, represented by *buzzatii* cluster species; 6) the *mojavensis* cluster; and 7) the *serido* sibling set, including *D. antonietae, D. borborema* and *D. koepferae*, three species that breed mainly on columnar cacti, as opposed to the prickly pear dweller *D. buzzatii*.

#### 2.7.2 Validation of candidate TRGs

Validation of TRGs involved the search of divergent homologs for each candidate TRG in a focal lineage. Then, we classified each candidate orthogroup as a divergent TRG (has distant homologs) or as a validated TRG (does not have distant homologs). The ancestral branch (root) was excluded from this step due to the lack of outgroups for the genus *Drosophila*. To this end, two filters were applied to classify candidate TRGs as well as to find differences with potential candidate TRGs. Firstly, clustered proteins of each candidate orthogroup detected in a focal lineage (branch) were mapped to the genomes of the outgroup species using TBLASTN with *e-value* and coverage match cutoffs of 1×10^-3^ and 50%, respectively. In this mapping, we employed a more relaxed *e-value* threshold than in the ortholog identification method in search of homologous genes which were already annotated but diverged beyond the recognition of sequence similarity methods. We classified a candidate orthogroup as a divergent TRG if every protein member had at least one hit to a genome region where a gene was annotated in an outgroup. A similar criterion was applied in BLASTP comparisons against the non-redundant reference proteomes release 1444 database available at RefSeq (Pruitt et al., 2007). This conservative filter was used to detect distant relationships given the substantial divergence between species included in this study. All candidate orthogroups with no hits to outgroups were classified as validated TRGs.

We compared the distribution of AED score and protein length for the validated and divergent candidate TRGs in the *buzzatii* species cluster branch against the set of toolkit genes, i.e. the sets of conserved genes across the different lineages in the species tree. We also evaluated the normal distribution in AED scores of divergent TRGs and validated TRGs using Shapiro-Wilk’s test (toolkit genes were not analyzed because of a sample size greater than 5,000).

#### 2.7.3 Molecular evolution of TRGs

We investigated patterns of molecular evolution of all candidate TRGs considering only one-to-one orthogroups. First, fasta format files were generated with the amino acid sequences of ortholog genes, and MAFFT was employed for sequence alignment using the parameters “--unalignlevel 0.1 --leavegappyregion --globalpair --maxiterate 1000”. Second, nucleotide sequence alignments corresponding to coding regions were built following the same steps as for amino acid alignments. Codon alignments were subsequently generated for each orthogroup by applying PAL2NAL ver. 14 (Suyama et al., 2006) to each pair of amino acid and nucleotide sequence alignments. Amino acid alignments were additionally refined with trimAl to infer orthogroup phylogenetic trees using IQ-TREE with default parameters. Non-synonymous to synonymous substitution rates (dN/dS) ratio (ω) across ortholog codon sequences were calculated in search of positive selection using the program codeml of the PAML ver. 4.9 package (Yang 1997, 2007). The BioPython PAML module was used to create control files and to test the fit of different codon models to the observed data. In each case, the control files were configured to employ the corresponding ortholog gene tree and codon alignment. In this way, model M0 was first fitted to the data to estimate one single average ω for each orthogroup and to obtain branch lengths to be used as initial values for more complex models. The models M7 with 10 omega site classes not allowing positively selected sites and M8 with an extra class constrained to have ω ≥ 1 were fitted to the data to estimate model log-likelihood (L). Model M8a in which the extra class in M8 is fixed to 1 (ω = 1) was used as an alternative null hypothesis to avoid false positives. L values of the models tested were compared using likelihood ratio tests (LTR) with *α* set to 0.05. The sequential Bonferroni correction was then applied to correct *α* for multiple testing. Hence, the LTR statistic was computed among models employing the following equation: 2 x (L1 - L0), where L1 and L0 are the log-likelihood values of the different hypotheses (models) tested. The LTR values were contrasted against the chi-square (𝜒2) distribution considering the degrees of freedom between models and the *α* value. Simpler models were rejected in favor of more complex ones in each comparison when the LTR value was greater than the 𝜒2 value.

#### 2.7.4 TRG functional prediction

Functional annotation of TRGs identified in each branch of the species tree was made with eggNOG-mapper (Huerta-Cepas et al., 2017) to predict orthology (one2one) with eukaryotic proteins in the eggNOG database (Huerta-Cepas et al., 2019). Gene Ontology (GO) annotations were only transferred to candidate TRGs if all orthologs had the same match. Annotated TRGs in each branch of the species tree were used to perform an enrichment analysis (FDR < 0.05) by testing the overrepresentation of annotated GO terms for the sets of TRGs against the functional background, i.e. the GO terms annotated in *D. melanogaster* proteins for orthogroups in the tree root. Revigo (Schlicker et al., 2006; Supek et al., 2011) was employed to reduce the redundancy of enriched GOs using the simRel semantic similarity score (Schlicker et al., 2006) and the *D. melanogaster* UniProt reference dataset of GOs was used to obtain specificity (frequencies) of all recovered terms.

## 3. RESULTS AND DISCUSSION

### 3.1 Sequencing and genome assembly

We obtained coverage values ranging from 110x to 136x among samples. *Drosophila koepferae* A had considerably higher coverage (262X) given additional Illumina reads obtained previously in our lab (De Panis et al., 2016). These values were calculated on the basis of the genome size of ~160Mb estimated for *D. buzzatii* (Guillén et al., 2014). For all species, genomic DNA was assembled into less than 642 contigs and genome size varied between 166 and 191 Mb. These numbers are similar to those obtained in other genomic projects involving species of the *repleta* group (Table 1 and Table S1).

**Table 1.** Contiguity statistics for 13 genome assemblies.

| Assembly | Dald | Dari | Dato | Dbrb | Dbuz | Dhyd | DkoeA | DkoeB | Dmel | Dmoj | Dnav | Dmex | Dvir |
| --- | --- | --- | --- | --- | --- | --- | --- | --- | --- | --- | --- | --- | --- |
| Total length (Mb) | 190.65 | 141.39 | 166.31 | 191.23 | 161.49 | 153.74 | 180.74 | 171.78 | 143.74 | 193.82 | 147.36 | 163.50 | 20.60 |
| # contigs/scaffolds | 2,620 | 3,178 | 642 | 364 | 826 | 217 | 609 | 246 | 1,870 | 6,817 | 13,813 | 4,037 | 13,415 |
| Largest seq (Mb) | 4.1 | 29.98 | 32.47 | 27 | 16.30 | 15.87 | 10.93 | 19.20 | 32.08 | 34.15 | 3.64 | 9.86 | 25.23 |
| GC (%) | 40.04 | 40.34 | 39.5 | 38.93 | 38.48 | 39.45 | 40.23 | 38.32 | 42.01 | 39.48 | 39.78 | 40.69 | 39.99 |
| N50 (Mb) | 1.03 | 26.54 | 5.064 | 11.35 | 1.38 | 3.37 | 2.28 | 16.45 | 25.29 | 24.76 | 0.39 | 3.56 | 10.16 |
| N75 (Mb) | 0.30 | 23.95 | 0.26 | 2.96 | 0.53 | 1.77 | 0.96 | 5.97 | 23.54 | 3.41 | 0.05 | 1.90 | 2.01 |
| L50 | 53 | 3 | 7 | 6 | 30 | 9 | 21 | 5 | 3 | 4 | 80 | 13 | 6 |
| L75 | 139 | 4 | 52 | 15 | 76 | 24 | 50 | 11 | 4 | 8 | 370 | 30 | 20 |

The official FlyBase abbreviations for *Drosophila* species names are shown. Dald: *D. aldrichi*; Dari: *D. arizonae*; Dato: *D. antonietae*; Dbrb: *D. borborema*; Dbuz: *D. buzzatii*; Dhyd: *D. hydei*; Dkoe: *D. koepferae*; Dmel: *D. melanogaster*; Dmex: *D. mercatorum*; Dmoj: *D. mojavensis*; Dnav: *D. navojoa*; Dvir: *D. virilis*.

We compared the newly assembled genomes with nine sequenced *Drosophila* species and employed the genomic data of all species to perform a comparative genomic analysis. N50 and L50 statistics varied from 2.3 to 16.4 Mb and from 5 to 21 sequences, respectively, for the assemblies reported in this study (Table 1). These differences are probably related to variation in PacBio read length distribution among samples. Even though PacBio sequencing throughput was comparable among samples (5-5.5 Gb), the mean read length was, at least, 36% lower for *D. koepferae* A, which had the lowest N50 score (Table 1), than the other genomes (see more details in Table S2). The impact of read length distribution differences on genome assembly can also be observed in the length variation of the largest scaffold across samples, which ranged from ~11 Mb in *D. koepferae* A to ~32 Mb in *D. antonietae*.

Contiguity can also be influenced by several factors such as heterozygosity, sequencing depth, and repeat content (Yandell & Ence, 2012). For example, diploid or even polyploid genomes present further complexity than prokaryotic genomes, challenging assemblers to resolve regions enriched with paralogs or determine regions where the assembly yielded separated contigs/scaffolds caused by high heterozygosity. Thus, sequencing inbred lines helps to obtain deeper coverage and to increase assembly sensitivity given the presence of mostly single allelic positions along a potential haploid genome (Huang et al., 2017; Zhang et al., 2020). In this sense, our hybrid assembly approach combined short and long sequencing reads. This probably aided in the resolution of large repeats, allowing the assembler to span these complex regions with long PacBio reads (Rhoads & Au, 2015), as well as to balance the high error rate by mapping the Illumina reads (Walker et al., 2014).

The mean number of unknown nucleotides (N-positions) per 100 kb was zero in the new genomes, which is consistent with our protocol assembly that did not address scaffolding steps after merging only aligned sequences between the two initial assemblies. These results contrast with genome assemblies obtained in the other species, which had 4-9% of the genome with non-resolved positions since they were obtained using protocols that try to reach full chromosome scaffolds (e.g. Guillén et al. 2014). Chromosome-level assemblies generally have higher contiguity due to the presence of scaffolds composed of contigs joined by N-positions (after scaffolding). The highest N50 scores among the 13 genome assemblies compared herein were obtained in *D. arizonae* (26.6 Mb), *D. melanogaster* (25.3 Mb), and *D. mojavensis* (24.8 Mb), with contiguities at least 8 Mb higher than that of *D. koepferae*, which had the highest N50 score among the newly reported genomes. However, this statistic decreases drastically in most of the other genomes if we consider contigs instead of scaffolds. To further analyze this issue, we compared the scaffold and contig N50 scores after splitting scaffolds in contigs by removing N-positions in each of the 13 genome assemblies (Figure 1). *Drosophila melanogaster*, on one hand, and the remaining outgroup species, on the other hand, had better and worse contiguity values than the newly genomes reported herein, respectively (Table 1). As expected, the use of long reads had a great impact on contiguity, as it is mostly reflected in the fact that our assemblies contained half of the genome in less than 21 sequences while most other genomes required at least 200 contigs (Figure 1 and Table S3).

**Figure 1.**
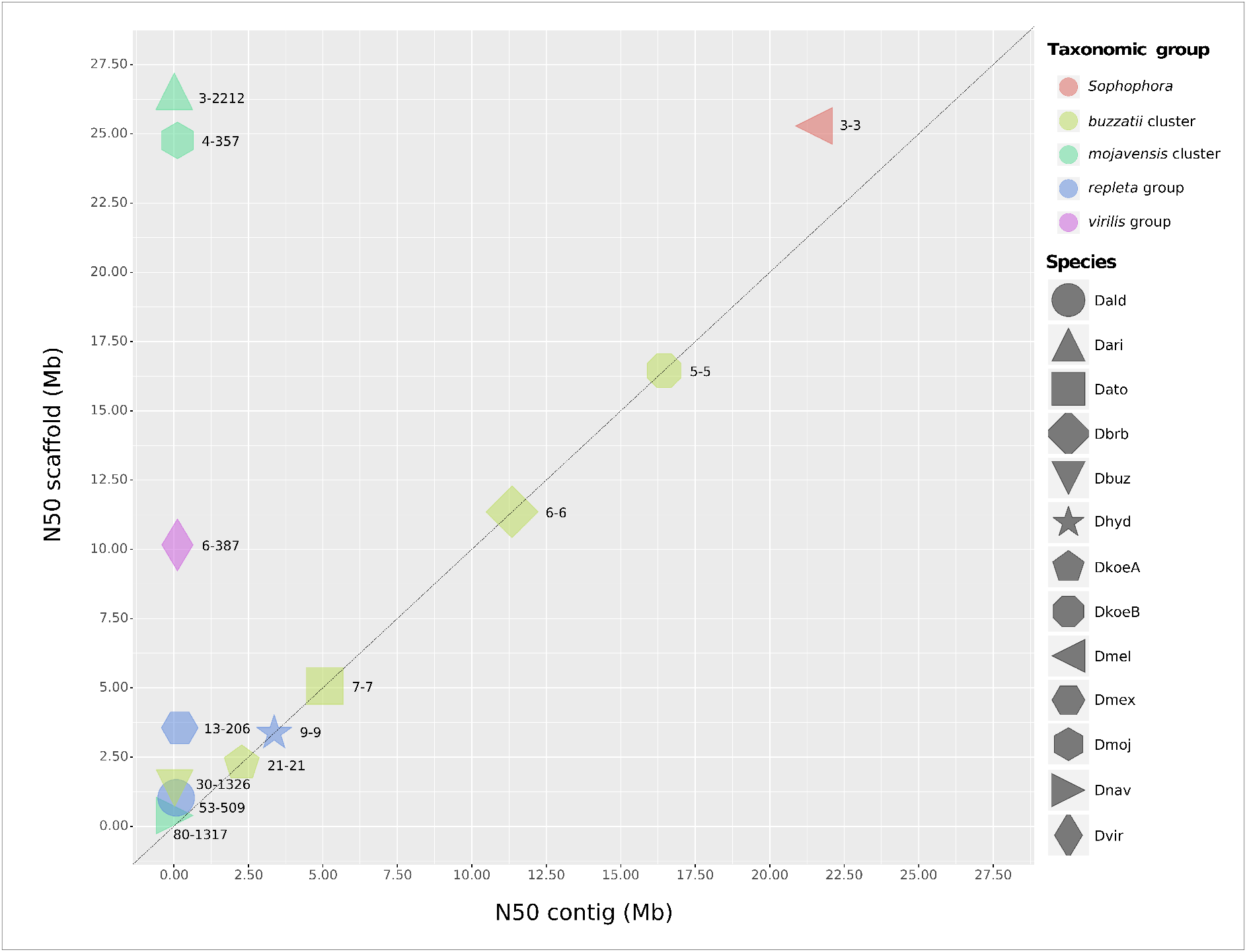
Scaffold and contig N50 score comparison. Each species’ genome assembly is represented by a different geometric shape. The color of each shape indicates the taxonomic group of the corresponding species. The diagonal dashed line shows the perfect correlation between both scores. Values separated by a hyphen next to each genome assembly indicate the scaffold and contig L50 score.

The evaluation of gene representation is frequently employed to assess genome assembly quality. We evaluated genome completeness using dipteran BUSCO groups, i.e. single-copy orthologs (Simão et al., 2015). These searches yielded completeness values above 96.5% in all new assemblies (see Figure S2 and Table S3). Furthermore, the number of fragmented genes was almost always lower than 20 with slight differences among assemblies. The numbers of missing genes were slightly higher in *D. antonietae* (94) and *D. koepferae* A (81) than in *D. borborema* (28) and *D. koepferae* B (32). Similar estimates were also obtained for *D. aldrichi, D. arizonae, D. buzzatii, D. navojoa,* and *D. virilis* (Miller et al., 2018; Jauhal & Newcomb, 2021).

It has been recently shown that there is a positive correlation between N50 and complete BUSCO scores among eukaryotic genomes and that haploid or highly homozygous (our case) genomes are expected to present best single-copy completeness scores since assemblers do not have to overcome the obstacle of distinguishing allelic variants from duplications (Jauhal & Newcomb, 2021). Therefore, assessing assembly quality by complementing contiguity and completeness statistics allowed us to validate our sequencing protocol and to demonstrate that the genomes reported herein are of comparable quality to other *Drosophila* species.

### 3.2 Repeat content variation among genomes

We constructed species-specific repeat libraries aiming to mask genomes before annotation as well as to assess repeat content. We found that ~19-21% of the four new genomes are composed of repetitive sequences, with *D. borborema* showing the highest proportion of repeats (21.12%) (Table S4). In all cases, more than half of repeat content (10.25-12.15%) consists of interspersed sequences such as transposable elements (TEs), followed by simple repeats (7.25-8.16%) and low complexity sequences (~1%). These results are similar to those obtained in *D. buzzatii*, though data for other repetitive elements were not reported (Rius et al. 2016). Between 8.22 and 10.62% of total repeats were classified as unknown by RepeatMasker. To validate these results, we used the reference *Drosophila* repeat library, available in the Dfam database (Hubley et al., 2016), to mask each new genome. The results were very similar to those obtained with the first approach, although the percentages of interspersed elements were slightly lower (1-2%). In addition, the number of repeat sequences classified as unknown was considerably reduced to 0.15-0.19% (Table S4), as the Dfam repeats are mostly annotated. The high percentage of unclassified sequences using the first approach could be due to the use of only representative sequences (not all individual repeat sequences) to mask each genome (see Methods).

### 3.3 Genome annotation assessment

The number of annotated genes varied widely among assemblies. On one hand, the numbers of genes detected in *D. buzzatii* (13038) and *D. koepferae* B (14852) are in agreement with reports in species of the *mojavensis* cluster (Table S5). On the other hand, more than 17,500 genes were annotated for *D. antonietae, D. borborema*, and *D. koepferae* A, which is comparable to the number reported for *D. melanogaster* (Table S5). Average gene length fluctuated from 3,411.6 to 4,111.6 bp in *buzzatii* cluster genomes. These values are smaller than the average gene length of 6,117.9 bp calculated for the other genomes included in this study. This dissimilarity is probably related to a lower amount of data and the types of tissues covered in the transcriptomic evidence used for annotation in this study. The number of species-specific transcripts mapped to each new genome was limited, and assembled by employing only between 29 and 45.5 million RNA-seq reads that were obtained from adult male whole body and reproductive accessory glands and adult female whole body (see Methods). The reannotation of the *D. buzzatii* genome using additional data resulted in an increased average gene length. The first annotation of this species’ genome was based on almost 300 million reads encompassing four developmental stages (Guillén et al., 2014), resulting in 13,657 annotated genes with an average length of 3,107.1 bp. After re-annotating the *D. buzzatii* genome using two additional transcriptomic datasets (120 million reads), the mean gene length increased by ~1 kb (4,111.6) and the number of genes was slightly reduced (13,038 genes).

All genome annotations had at least 95% of gene models with AED values of 0.5 or better among annotations (see Figure S3). In particular, the re-annotation of *D. buzzatii* yielded slightly better results than the remaining *de novo* annotations. Nevertheless, we could not compare this metric with *Drosophila* reference species or even with the first version of *D. buzzatii* because of the lack of AED values associated with their annotated gene models. However, these distributions reached the standard threshold used to determine a good annotation quality in model species (Eilbeck et al., 2009; Holt & Yandell, 2011; Cheng et al., 2017; Baxevanis et al., 2020).

### 3.4 Phylogenomic analyses

Phylogenetic relationships were inferred by employing the protein alignment of 1,866 BUSCO groups. Although all genomes showed completeness scores above 96% for 3,285 dipteran genes (see Figure S2), only 57% of the alignments, corresponding to shared singlecopy orthologs among all species, were concatenated into the matrix used for phylogenetic inference. This reduction in the number of BUSCO groups was mostly caused by missing and fragmented genes, as well as by the presence of duplicated genes.

The species tree was inferred using 1,312,819 amino acid sites and the topology retrieved (Figure 2), where all nodes had the highest support values (100%), is consistent with expectations for the taxonomic groups included (Text S1). All in all, our phylogenomic approach is in agreement with pre-genomic studies, which reported the monophyly of the *repleta* group and the relationships among species of the six subgroups (three of which are included in this report: *hydei, mercatorum*, and *mulleri* subgroups) (Oliveira et al., 2012). Furthermore, the *buzzatii* cluster, representative of the *buzzatii* complex in this study, appears as the sister clade of the *mulleri* complex (i.e. the *mojavensis* cluster +*D. aldrichi*). These two complexes, which are part of the *mulleri* subgroup, comprise the sister branch of *D. mercatorum* (*mercatorum* subgroup). As shown by Oliveira et al. (2012) and Suvorov et al. (2021), *D. hydei* represents the outermost branch of the *repleta* group clade. However, the positions of *D. hydei* and *D. mercatorum* in our tree are at odds with recent reports showing *D. mercarotum* as the most basal species of the clade (Li et al., 2021; Rane et al., 2019). Relationships within the *buzzatii* cluster in our tree are inconsistent with previous studies (Hurtado et al., 2019; Moreyra et al., 2019), particularly for the trio *D. antonietae, D. borborema,* and *D. koepferae*, which are representatives of the main lineages defined on grounds of fixed chromosomal inversions in the *serido* sibling set (Ruiz et al., 2000; Hasson et al., 2019).

**Figure 2.**
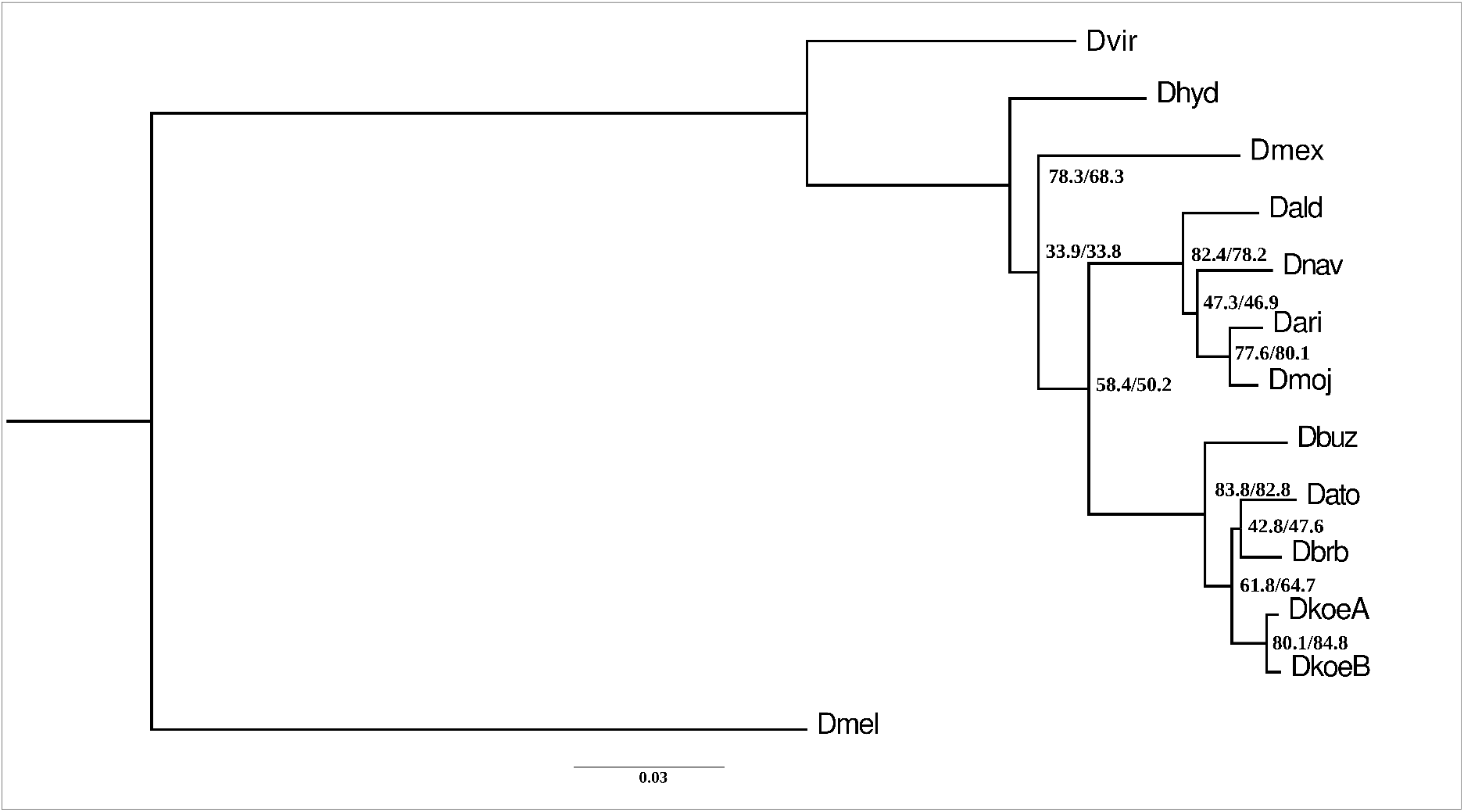
Phylogenetic tree inferred using dipteran BUSCO groups in Maximum Likelihood searches. Bootstrap values were equal to 100% for all nodes (not shown). Gene- and site-concordance factors values (separated by slashes) are shown for each node.

We also explored relationships by means of a gene/locus tree approach. In general, gCF and sCF values were similar in all nodes (see Figure 2 and Table S6), indicating that any difference between these factors and the results of the bootstrap analysis was likely caused by genuinely discordant signals in the gene trees rather than stochastic errors from limited information as short branches (Minh et al., 2020). Moreover, in four clades the gCF and sCF values were considerably different from bootstrap supports. Firstly, the clade formed by *D. mercatorum* and the *D. mulleri* subgroup had the lowest values, with only 33.9% of the gene trees supporting the species tree topology. This low value may be related to the short length of this specific branch and to genuinely contradictory signals in gene trees likely generated by incomplete lineage sorting (ILS) (Degnan & Rosenberg, 2006). We also looked at the percentage of genes that support alternative hypotheses for each specific branch in the species tree, the gene discordance factors (gDF) (Minh et al., 2020). Firstly, we found that 25.5% (gDF1) and 17.5% (gDF2) of the genes supported alternative relationships between *D. hydei, D. mercatorum*, and the *D. mulleri* subgroup, and that the remaining 23.1% is in discordance due to paraphyly (gDFp). Secondly, 58.4% of the gene trees agreed with the sister relationship between the *buzzatii* and *mulleri* complexes. The gDF1 and gDF2 were lower than 7% in this branch, but the gDFp was considerably higher, reaching almost 30%. Thirdly, the placement of *D. navojoa* as the first branch splitting off in the *mojavensis* cluster was supported by less than 50% of the genes and sites, whereas alternative relationships and genes in discordance represented 40% and 14%, respectively. Lastly, the *buzzatii* cluster was well-supported by both factors, ~83 and 84%, and the position of *D. buzzatii* as the sister species of the *serido* sibling set was supported by approximately two thirds of the genes and sites (~62-65%). In this sense, the basal position of *D. buzzatii* is in agreement with previous studies (Rodriguez-Trelles et al., 2000; Oliveira et al., 2012; Hurtado et al., 2019) but differs from studies based on mitochondrial markers and mitogenomic data (Manfrin et al., 2001; Moreyra et al., 2019). In turn, the subclade containing *D. antonietae* and *D. borborema* was supported by less than 50% of genes and sites, while 1067 genes (57%) retrieved alternative hypotheses. Such inconsistent relationships within the *serido* sibling set may be caused by ILS and/or interspecific gene flow, as it was already proposed in a phylogenomic study based on transcriptomic data (Hurtado et al., 2019). These authors reported a large discordance among gene trees and suggested that the pattern of divergence in this trio represents a hard polytomy. In agreement with these results, the *serido* sibling set clade had 15.6% of the genes supporting two different resolutions and 22.6% in discordance due to paraphyly.

### 3.5 Divergence times

The node age calibration approach relied on the species tree obtained with the BUSCO matrix (Figure 2), estimating only branch lengths and node ages, whereas the mutation rate calibration approach employed phylogenetic searches along with the estimation of node ages. Divergence time estimates obtained with the two methods are shown in Figure 3. The mutation rate calibrated method resulted in a topology slightly different from the species tree shown in Figure 2 since *D. hydei* and *D. mercatorum* appear as sister species. This difference in topology may be due to differences in the datasets used in each approach. The mutation rate-based method is only applicable to neutral sites and, thus, we selected and concatenated into the matrix 29,923 four-fold degenerate third-codon sites (4FDS) extracted from 151 genes with low codon bias (see Methods). The species tree, instead, was based on amino acid sequences of 1,866 BUSCO groups.

**Figure 3.**
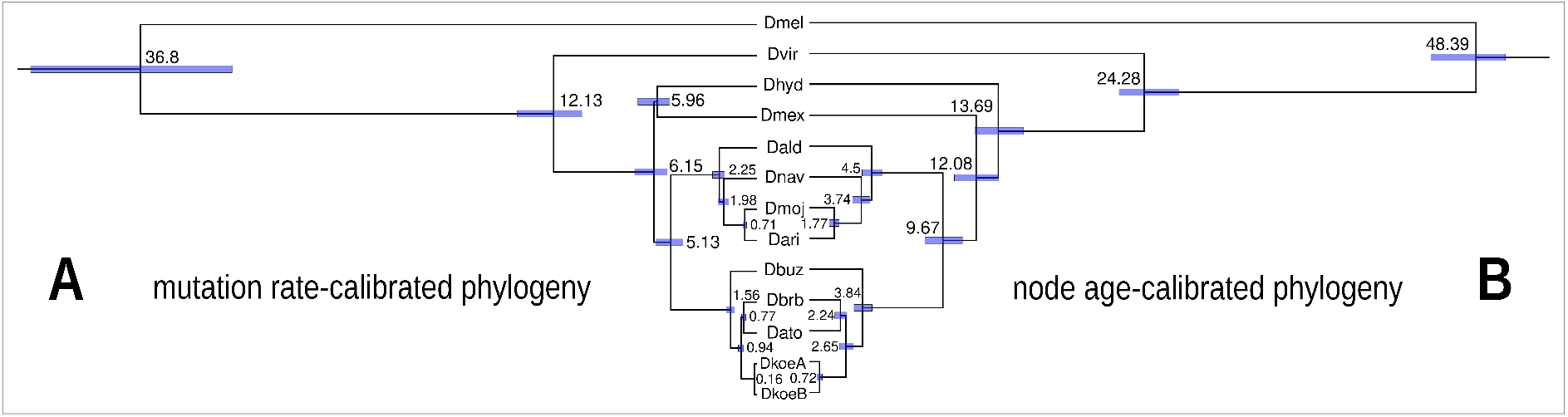
Divergence times and topologies recovered with two approaches differing in the priors used to calibrate the phylogeny. The numbers on each node are the time estimates. Blue bars represent their 95% confidence intervals of estimates. The topology on the left (A) represents the time estimates obtained using a mutation rate to calibrate the phylogeny and the topology on the right shows divergence times calculated using previously reported node ages as calibration prior.

The estimates obtained with the node-calibration method suggest that the *virilis* and *repleta* groups shared their last common ancestor about 24.3 Mya and that the *mulleri* and *buzzatii* complexes diverged 9.7 Mya, somewhat later than the estimates reported in Oliveira et al. (2012) using the same approach. The first split separating *D. navojoa* from the clade *D. arizonae-D. mojavensis* within the *mojavensis* cluster is suggested to have occurred more recently (3.7 Mya) than previously reported (Machado et al., 2007; Oliveira et al., 2012; Sanchez-Flores et al., 2016) but is in line with estimates based on Alcohol dehydrogenase (ADH) (Matzkin & Eanes, 2003). The time of separation between *D. arizonae* and *D. mojavensis* was very close (~1.8 Mya) to estimates obtained in the above-mentioned studies. Within the *buzzatii* cluster, the divergence of *D. buzzatii* from the ancestor of the *D. serido* sibling set was estimated to have occurred 3.8 Mya, whereas speciation events within the *D. serido* sibling set appeared very close in time (2.2 and 2.7 Mya), in accordance with the idea of a hard polytomy (Hurtado et al., 2019). The results obtained for the *buzzatii* cluster are similar, though slightly more recent, to those reported in (Oliveira et al., 2012). Lastly, divergence between Bolivian and Argentine *D. koepferae* using the node age constraint method is more than twice (0.7 Mya) the estimate obtained with mitogenomes (0.3 Mya) (Moreyra et al., 2019).

The comparison of divergence times obtained with the two approaches revealed discrepancies at all nodes (see Figure 3 and Table S7). Overall, the mutation rate-calibrated phylogeny yielded divergence times that are more recent than those based on the node age-calibrated phylogeny, a pattern that is in agreement with results from previous studies in *Drosophila* (e.g. Obbard et al. 2012; Sanchez-Flores et al., 2016; Hurtado et al. 2019; Suvorov et al. 2021). For instance, divergence time estimates obtained with the mutation rate method are in agreement with those reported by Obbard et al. (2012) using the same approach for both the species tree root (*Drosophila-Sophophora* split, ~37 Mya) and for the *virilis-repleta* radiation (~12 Mya). With this method, the origin of the *repleta* group was estimated at 6.2 Mya, which is at least twice more recent than the estimates reported in Oliveira et al. (2012) and herein using the node age calibration approach (16.3 and 13.7 Mya, respectively). This pattern repeated more or less itself throughout the phylogeny (Table S7).

In a recent study, Hurtado et al. (2019) estimated divergence times within the *buzzatii* cluster based on transcriptomic data using the same mutation rate approach to calibrate the clock. Accordingly, our estimate of divergence time between *D. buzzatii* and the ancestor of the *serido* sibling set was very similar to that obtained by those authors (1.6 Mya), suggesting that the diversification of the *buzzatii* cluster happened in the Pleistocene. Nevertheless, within the *serido* sibling set our estimates were slightly older (0.9 and 0.8 Mya) than those reported in Hurtado et al. (0.4 and 0.5 Mya). In both studies, however, speciation events within the *serido* sibling set appear to have occurred in close proximity to each other, again supporting the hypothesis of a hard polytomy for this trio (Hurtado et al., 2019).

The host specificity of most *repleta* group species suggests that their evolution should be synchronized with the evolution of cacti. It has been estimated that the Cactaceae family originated about 32.1 Mya and that its major diversification took place in the Miocene following the expansion of the New World’s arid and semi-arid lands 15-10 Mya (Hernández-Hernández et al., 2014). Our divergence time estimates based on node-age calibration placed the diversification of the *repleta* group (13.7 Mya) within the same time frame. Previous studies suggested that both cacti and cactophilic flies of the *repleta* group originated in central western South America (e.g. Oliveira et al., 2012; Hernández-Hernández et al., 2014). This synchronization gives support to the node age calibration approach, as the mutation rate approach suggests a delayed diversification of cactophilic flies. It is interesting to note, though, that the genus *Opuntia*, the most commonly used host plant by cactophilic species of the *repleta* group and also the proposed “ancestral host” (Oliveira et al., 2012), originated relatively recently, between 7.5 and 3 Mya (Hernádez-Hernández et al., 2014), which overlaps with the mutation rate estimates for crown *repleta*.The implications are that either *Opuntia* was not the ancestral host or that the mutation rate approach is more accurate. Alternatively, other genera within the speciose Opuntioideae subfamily, which diversified about 10 Mya (Hernández-Hernández et al., 2014), may have served as ancestral host plants of cactophilic *repleta* flies. In fact, most of the cactus genera that are currently used as breeding substrates by cactophilic species seem to have originated within the last 7 million years, which suggests host shifts throughout the history of the *repleta* group.

### 3.6 Orphan gene evolution

The search for potential orthologs involved 175,173 genes among the 13 proteomes of the species included in our study and retrieved 16,448 orthogroups (OGs) distributed across all internal branches of the phylogeny (see Table S8). We further analyzed the species tree to identify candidate orphans and TRGs shared by all species in each focal lineage (internal clade) with no detectable homologs in outgroup species. These candidates may be novel genes (validated TRGs), genes lost in external branches (validated TRGs), or genes that have diverged widely from their homologs (divergent TRGs).

One caveat of this analysis is that the gene sets of the species outside the *buzzatii* cluster were retrieved from different bioinformatic sources such as NCBI (Pruitt et al., 2007), FlyBase (Thurmond et al., 2019), and individual species sequencing projects (see Table S1), and, thus, annotations were not generated following the same methodology. For instance, the genomes of *D. arizonae, D. hydei*, and *D. navojoa*, were annotated using an automatic pipeline, whereas *D. melanogaster* annotations are periodically updated and manually curated. In addition, some annotation methods are fully predictive (Salamov & Solovyev, 2000; Aggarwal & Ramaswamy, 2002; Korf, 2004; Stanke et al., 2004), while others incorporate the information of RNA-seq reads and known protein mapping (Cantarel et al., 2008; Holt & Yandell, 2011; Campbell et al., 2014; Hoff et al., 2016; Tatusova et al., 2016; Thibaud-Nissen et al., 2016) as guides in the construction of gene models. Thus, differences between methodologies may generate a bias in the number of genes that can be recognized in each genome (Eilbeck et al., 2009; Weisman, 2021), even when comparing annotations obtained for a single species using different methodologies (Holt & Yandell, 2011; Casola, 2018; Zile et al., 2020). Therefore, there is an inherent error in comparative genomic analyses that must be taken into account when annotation heterogeneity exists among samples (Weisman, 2021). Considering this caveat, we report the annotations of four newly sequenced genomes of the *buzzatii* cluster and the re-annotation of the already sequenced genome of *D. buzzatii* (Guillén et al., 2014) using the same protocol to reduce biases.

The ancestral branch (root) had 6,941 OGs (Figure 4.A), though it is possible that additional OGs lost in some species were not considered in this count. We focused on the validation of candidate TRGs in the ancestral and internal lineages of the subgenus *Drosophila*, which was represented by species of the *virilis-repleta* radiation. Divergent TRGs were not considered as novel genes because these genes diverged from the respective presumptive homologs in the ancestor of the corresponding focal lineage. However, divergent TRGs have been conserved in specific clades after divergence from the preexisting homologs, suggesting that they may be of relevance in the evolution of adaptive traits in these lineages (Domazet-Loso & Tautz, 2003; Khalturin et al., 2009).

**Figure 4.**
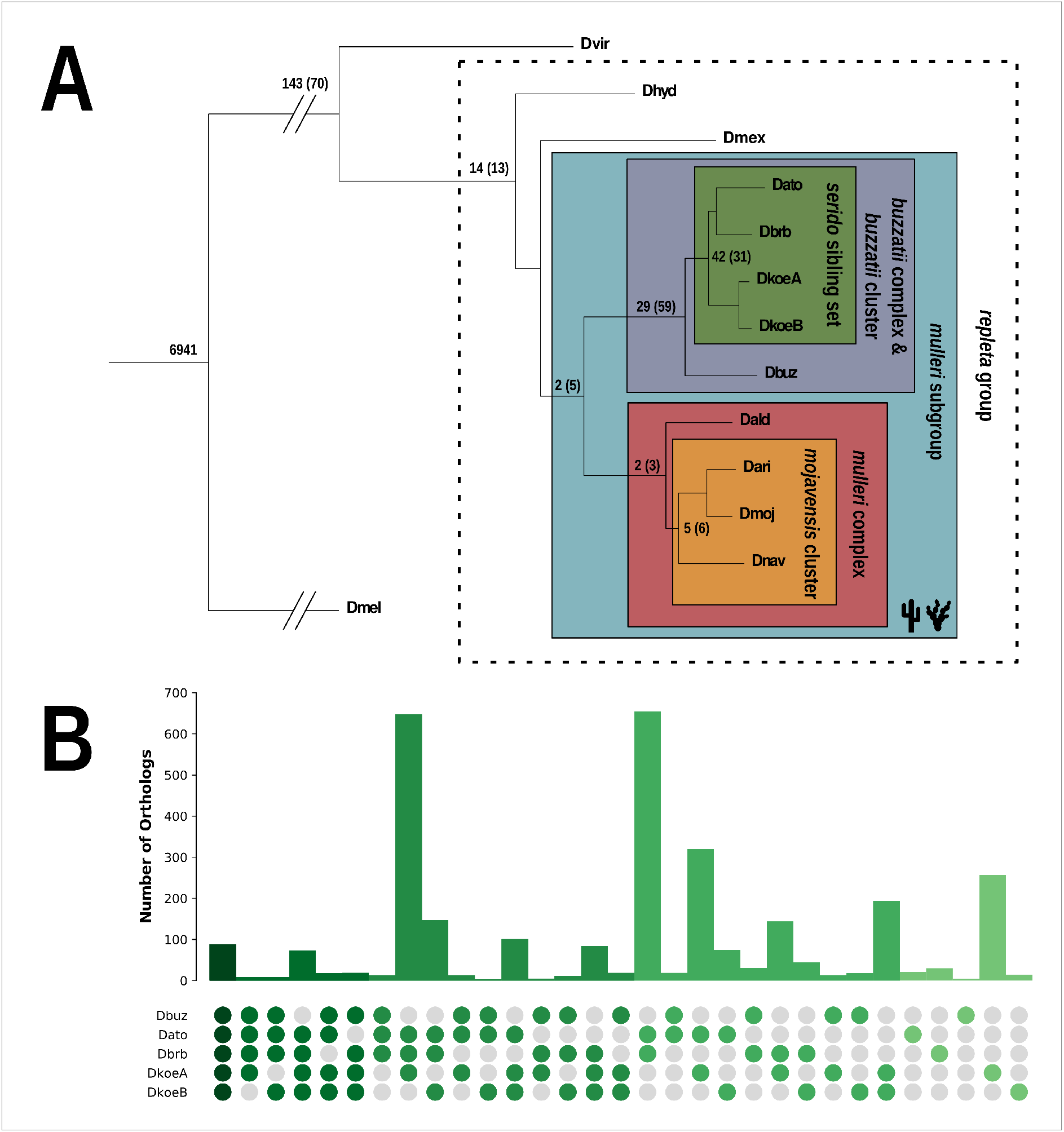
Identification of candidate TRGs. (A) Numbers of orthogroups (root) and TRGs in each branch of the species tree. The dashed line confines the *repleta* group, and the cactophilic mulleri subgroup and its internal lineages are framed in different colors. The values in each branch represent the number of validated and divergent (in parentheses) candidate TRGs. (B) Incomplete TRGs identified for each species combination within the *buzzatii* cluster. Each row of circles represents a single species and species combinations are shown in columns of colored circles. The color intensity represents the number of species included in each combination (from the darkest green for five species to the lightest green for one). The bars above each combination indicate the number of incomplete TRGs identified.

We found 143 validated TRGs out of 213 candidates in the subgenus *Drosophila* and 14 out of 27 in the *repleta* group (Figure 4.A). In the *mulleri* subgroup, which includes the *buzzatii* and the *mulleri* complexes (all species are cactus specialists), we identified seven candidate TRGs, of which two were classified as validated and five as divergent. We detected two validated and three divergent TRGs in the *mulleri* complex and out of a total of 11 candidate TRGs in the *mojavensis* cluster, five were validated and six were divergent. In addition, 29 validated and 59 divergent TRGs were detected in the *buzzatii* cluster and 42 validated and 31 divergent candidates in the *serido* sibling set. Finally, 5 genes with no detectable homology in other species (after validation) were shared between the generalists *D. hydei* and *D. mercatorum*. Although these two species do not form a clade in the species tree (but see the topology obtained in the mutation rate-calibrated phylogeny in Figure 3), it may be considered that those genes appeared either in the ancestral branch of the *repleta* group (later lost in the lineage leading to the *mulleri* subgroup) or in the common ancestor of *D. mercatorum* and *D. hydei*.

We deepened our search for TRGs in the five genomes of the *buzzatii* cluster and found genes that were present in some but not all members of the *buzzatii* cluster or *serido* sibling set. A total of 2,937 OGs were found in 29 species set combinations (considering orphans) (Figure 4.B and Table S9). Since some of these genes cannot be considered as restricted to a monophyletic clade (i.e. it is present in some but not all species of the clade), we named them incomplete TRGs; even in the case of *D. antonietae* and *D. borborema* that composed a clade in the species tree because of the unclear relationship with *D. koepferae* (Hurtado et al., 2019; Moreyra et al., 2019). Each one of the species of the *serido* sibling set shared less than 32 incomplete TRGs with *D. buzzatii.* Further, pairwise comparisons within the *serido* sibling set showed that *D. antonietae* and *D. borborema,* the more closely related species (see Figure 2), shared 654 incomplete TRGs and that 648 and 147 TRGs were found after including *D. koepferae* A or *D. koepferae* B to this pair, respectively. In addition, we separately revised the number of orthologs between *D. antonietae* and *D. borborema* with both *D. koepferae* strains. We found that *D. antonietae* shared 320 incomplete TRGs with *D. koepferae* A, 74 with *D. koepferae* B, and 101 with both strains. The number of orthologs shared between *D. borborema* and *D. koepferae* was slightly lower, 144 with the Argentine, 44 with the Bolivian, and 84 with both lines. Lastly, we detected species-specific orphans in all species: 21 in *D. antonietae,* 4 in *D. buzzatii*, 30 in *D. borborema*, and 194 in *D. koepferae* (both strains).

We dubbed incomplete TRGs the orthogroups for which a homolog could not be detected in one or more species either in the *buzzatii* cluster or the *serido* sibling set. Nevertheless, the absence of homologs in the genomic data of a given species may be the consequence of sequencing and/or assembly errors or miss-annotations. In cases where a TRG ortholog is indeed missing from the genome of one or more species, it is likely that the TRG was present in the ancestor of the *buzzatii* cluster (or *serido* sibling set) and was subsequently lost in one or more species or internal lineage. In this sense, it has been demonstrated that young genes (i.e. genes that emerged in a short branch) arise quickly and also have more chances to be lost (Tautz & Domazet-Lošo, 2011; Palmieri et al., 2014). Young genes tend to have relaxed selective constraints (Cai & Petrov, 2010) and therefore are more prone to gain indels and/or nonsense mutations, leading to pseudogenization (Palmieri et al., 2014). This may probably be the case for most of the non-spurious incomplete TRGs reported here, suggesting that they may not be novel genes with key adaptive roles but have not yet had enough time to get lost in all taxa.

We would like to remark that orphan and TRG candidates reported herein are working hypotheses. Further analyses are necessary to confirm if they are actual novelties or have diverged from distant homologs. For instance, a synteny-based approach aimed to search putative TRGs in conserved syntenic positions (Vakirlis et al., 2020; Zile et al., 2020), may help to confirm candidates that have originated by sequence divergence of ancestral genes.

We compared annotation accuracy (AED scores) and protein length distributions of the sets of validated TRGs (469 genes), divergent TRGs (366), and toolkit genes (51,155). Toolkit genes showed a lower mean AED score than both sets of TRGs (Figure S4), which had more similar AED values to each other. We could not test whether those differences in AED were significant as AED values departed from normality for both sets of candidate TRGs (*p-value* < 0.05 in all cases) and variance homogeneity among the three sets were rejected (Levene’s test), precluding the use of parametric and non-parametric tests. We also tested for normality and variance homogeneity of protein length distribution for each set of genes and, as for the AED score, normal distribution and homoscedasticity were rejected in all cases. However, the three sets of genes showed dissimilar distributions of protein lengths (Figure S5): divergent TRGs showed the lowest median value (158), followed by the validated TRGs (233.5) and toolkit genes (458). This is in line with previous reports showing that orphans and TRGs tend to be shorter (Lipman et al., 2002; Carvunis et al., 2012; Palmieri et al., 2014; Vakirlis et al., 2020) than other genes. Also, it has been shown that TRGs have low expression levels (Carvunis et al., 2012; Palmieri et al., 2014), offering a testable hypothesis for the TRGs identified herein.

### 3.7 Molecular evolution of TRGs

Our analyses showed that 51 out of 424 candidate TRGs distributed across all taxonomic groups in the species phylogeny (excluding the root) evolved under positive selection (see Figure 5 and Table S10). These TRGs with positively selected sites, of which 27 were validated and 23 were classified as divergent, were distributed across almost all analyzed lineages. Four positively-selected validated TRGs belong to the *virilis-repleta* radiation (subgenus *Drosophila*), 4 TRGs (1 validated TRG and 3 divergent TRGs) to the *repleta* group, 2 TRGs (1 validated and 1 divergent) to the North American *mulleri* complex, and 3 TRGs (2 validated and 1 divergent) to the *mojavensis* cluster. Our focal species had the largest numbers of TRGs evolving under positive selection: 11 validated and 15 divergent TRGs in the *buzzatii* cluster and 8 validated and 4 divergent TRGs in the *serido* sibling set.

**Figure 5.**
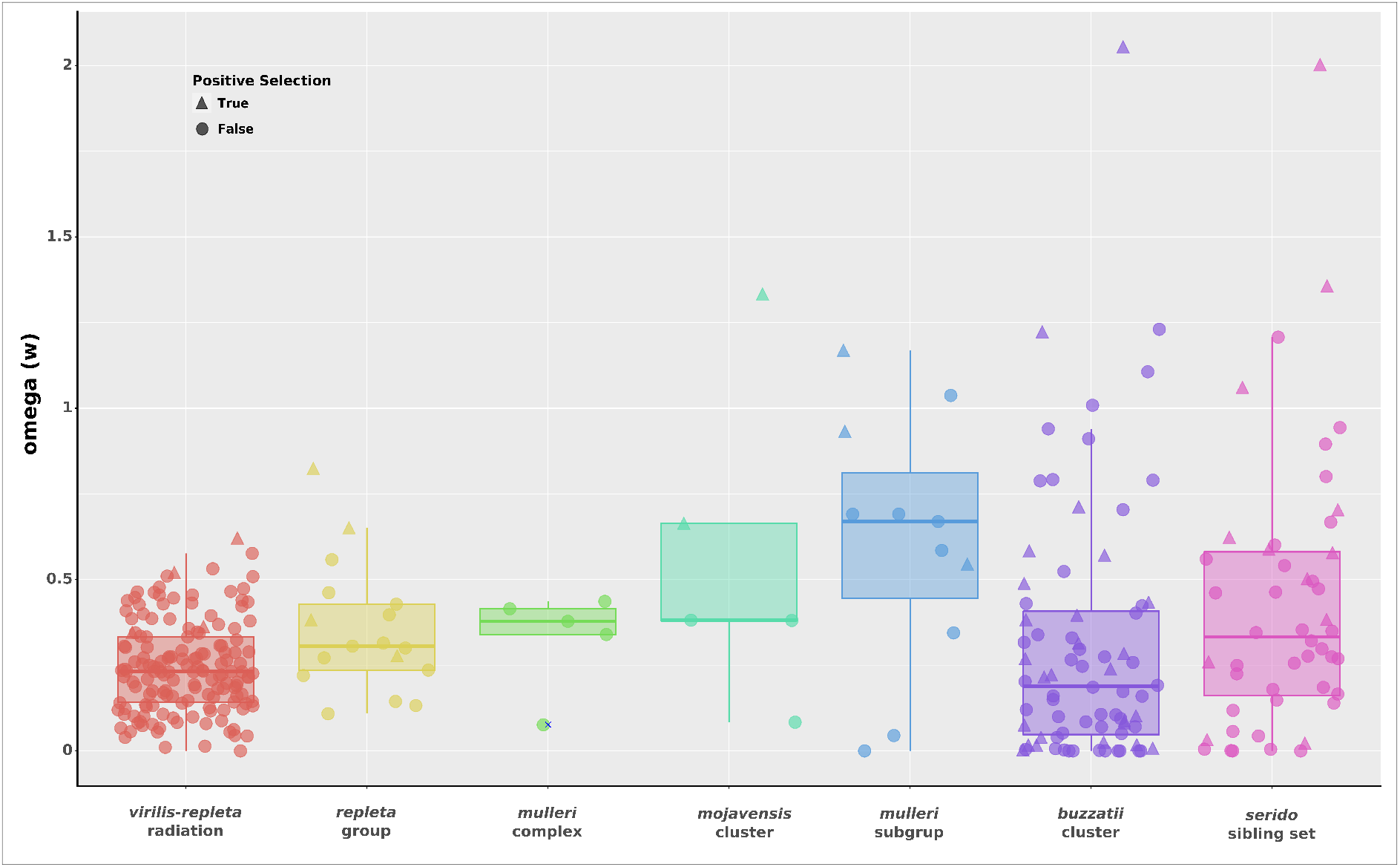
ω values for candidate TRGs in each branch of the species phylogeny. For each branch, significant TRGs for the positive selection test are represented by triangles and TRGs with non-significant results by circles.

### 3.8 TRG functional prediction

In these analyses, we focused mostly on the evolution of potential novel functions in the *repleta* group and internal lineages (see Table S11-12). Unfortunately, most TRGs belonging to the *repleta* group lacked annotated GOs, precluding functional enrichment testing. We detected only one functionally annotated TRG in the *mulleri* subgroup and one in the *mojavensis* cluster. The TRG of the *mulleri subgroup* had several annotated GO terms involved in cuticle development. Cuticle is the body outer layer that represents a barrier against pathogens and mechanical, physical, and chemical stresses (Moussian, 2010), and has been associated with functions involved in not only conferring more stability and water loss avoidance (Gibbs, 1998; Jaspers et al., 2014) but also working as a shield against xenobiotics (Agrawal et al., 2014; Kelkenberg et al., 2015). The *mojavensis* cluster annotated TRG appears to be involved in the transport of nitrogen compounds such as amides and peptides. In insects, this function is related to the excretion of nitrogenous waste (Weihrauch et al., 2012), which is key to adaptation to a xeric environment (Tasaki et al., 2017; Weihrauch & O’Donnell, 2021).

Functional prediction of TRGs in the *buzzatii* cluster (8) and in the *serido* sibling set (5) provides an enthralling picture of the genetic mechanisms in which they may be involved. Interestingly, some of the enriched GO terms were also recovered in comparative transcriptomic studies evaluating the effects of rearing on alternative cactus hosts and in the presence of phenylethylamine alkaloids isolated from the columnar cactus *T. terscheckii* in *D. buzzatii* and *D. koepferae* (De Panis et al., 2016, 2022 -submitted-). These TRGs are mainly involved in the regulation of stress and immune responses triggered by external stimuli such as chemicals, abiotic factors, pathogens, and chemotaxis. Delving into the broad range of compounds causing chemical stress, we found child GO terms related to the modulation of cellular responses to oxidative stress, toxic substances such as alkaloids and other drugs, nitrogen compounds, odorants, and food, as well as regulatory mechanisms of olfactory learning and glucose detection. Furthermore, two candidate TRGs are associated with cuticle development, as mentioned before for the *mulleri* subgroup. Other GO terms are related to several morphogenetic processes and the development of anatomical structures. Finally, two positively-selected divergent TRGs are linked to the regulation of nucleic acid-templated transcription.

For the TRGs identified in the *serido* sibling set, we obtained fewer enriched terms. The most common terms were related to the development of anatomical structures, though other significant GOs were associated with processes that regulate responses to abiotic stimulus and stress, as well as the metabolism of nitrogen compounds, cellular transportation, and secretion. In addition, two TRGs were associated with female mating behavior, one of which showed signs of positive selection.

Recently, Rane et al. (2019) reported a high frequency of gene gains in the branch of cactophilic species of the *repleta* group (*mulleri* subgroup) associated either with the acquisition of cacti and/or loss in the use of non-cactus hosts and with the spread into the American deserts. Thus, our present results are in line with Rane et al’s report, since a candidate novel gene found in the *mulleri* subgroup lineage is involved in cuticle development, which is the first line of defense against xenobiotics (Agrawal et al., 2014; Kelkenberg et al., 2015) and essential for desiccation and heat tolerance (Gibbs, 1998; Guo et al., 2022). Moreover, TRGs involved in adaptation to desiccation, heat, and chemicals emerged in the common ancestor of the *buzzatii* cluster. These novel genes are related to morphogenesis, development of cuticle and anatomical structures, which may provide resistance to extreme climates and tolerance to xenobiotics that flies may find in new hosts (Agrawal et al., 2014; Kelkenberg et al., 2015). Likewise, other TRGs detected in the *buzzatii* cluster are related to a broad range of GO terms such as responses to external stimuli and stress and the regulation of the immune system, which are deployed by larvae facing both chemical challenges (Kircher, 1982; Fogleman & Abril, 1990; Fogleman & Danielson, 2001) and pathogens present in cactus necroses (Hasson et al., 2019). New genes involved in the immune response against bacteria and yeasts would also play a key role in host plant adaptation. Previous studies reported that the proteasome system is implicated in the immune response (Mykles, 1999; Hoang et al., 2015), and may be regulated when insect larvae grow in alternative host plants (De Panis et al., 2016). Overall, based on these findings, we propose that genomic innovations in the *mulleri* subgroup and internal clades may have been driven by adaptation to both extreme climate conditions and to the use of cactus necroses.

Columnar cacti impose stressful conditions during larval development since many species contain toxic compounds such as alkaloids compared to the more benign environment offered by most *Opuntia* (De Panis et al., 2016). Previous research has shown that the chemical composition of columnar cacti negatively affects the development of *D. buzzatii* (an *Opuntia* breeder) but not the development of *D. koepferae* (Hasson et al., 2019). Notably, the set of GO terms enriched in the *serido* sibling set is consistent with the idea that functional innovations evolved as adaptations in the transition from prickly pears to chemically more complex hosts like columnar cacti. As explained above, these potential novel genes may have been of utmost relevance during adaptation to new hosts, allowing species to face the challenges posed by the chemically diverse resources. Lastly, one of two annotated TRGs related to the enriched GO term female mating behavior (a validated TRG), exhibited signals of positive selection. This result agrees with a recent report showing high rates of molecular evolution in genes involved in reproduction and mating behavior in populations of *D. mojavensis* using different cacti as breeding substrates (Allan & Matzkin, 2019).

## 4. CONCLUDING REMARKS

Comparative genomics in groups of species living in ecologically different contexts may provide clues regarding the genetic mechanisms that have been shaped by adaptation to cactus hosts. The newly reported genomic data allowed us to reconstruct the most complete phylogeny to date of the *buzzatii* cluster as well as to estimate the divergence times of these species in active cladogenesis. Also, based on the inferred phylogenetic relationships, we report sets of candidate orphans and TRGs in the internal branches of the subgenus *Drosophila*, which seems to have emerged either by sequence divergence from ancestral homologous and/or *de novo*. Regarding the *buzzatii* cluster and the *serido* sibling set, we functionally characterized the candidate TRGs shedding light on the instrumental mechanisms underlying ecological innovations. During the acquisition of cactophily in *Drosophila*, genomic changes likely drove the evolution of multiple performance traits. Many of them might be associated with tolerance to extreme climate conditions faced by cactophilic flies of the *repleta* group during expansion across the American deserts. In addition, other changes might have evolved as adaptations to ecologically and chemically different resources. Indeed, the specialization to cactus hosts is associated with the acquisition of new mechanisms involved in detoxification, water preservation, immune system response, development of anatomical structures and morphogenesis, behavior, reproduction, and metabolism. All in all, our study provides insights into the role of genomic changes that likely drove the evolution of novel traits associated with the acquisition of cacti as breeding and feeding sites, and further host specialization. However, genomes of other *mulleri* subgroup cactophiles (e.g. *longicornis* and *meridiana* complexes) and species that diverged earlier in the *repleta* group (e.g. *D. eremophila* complex) (Oliveira et al., 2012) are necessary to discern whether the evolution of novel genes involved in the functional processes identified herein are a common feature in this species group.

## ACKNOWLEDGMENTS

We thank Colline Jaworski for their advice in the adaptation of the genome assembly protocol. We are also grateful to Juan P. Hurtado, Diego De Panis, and Julián Mensch for sharing transcriptomic data employed in the genome annotation process. We express our gratitude to members of the Laboratorio de Evolución for helpful discussions and constructive criticisms. We acknowledge Hernán Dopazo for granting access to Biocodices computational facilities and for advice and encouragement in the earlier stages of the project. We are grateful to Centro de Computación de Alto Rendimiento (CeCAR) for allowing us to use computing facilities in the final stage of the study. We thank Martina Weil for their advice on the functional enrichment analysis. This work was supported with grants from Agencia Nacional de Promotión Científica y Técnica (ANPCyT), Consejo Nacional de Investigaciones Científicas y Técnicas (CONICET) and Universidad de Buenos Aires (UBA) awarded to EH. The funders had no role in the study design, data collection, analysis, decision to publish, or preparation of the manuscript.

## AUTHOR CONTRIBUTIONS

NNM and EH conceived and designed the study; NNM performed most and FCA part of data analyses; LMM and NF contributed materials and participated in different stages of the research; CA carried out part of the wet lab work; NNM designed and made all figures and tables; NNM and EH led the writing and FCA contributed to the discussions and writing of the manuscript; CA, NF and LMM revised previous drafts. All authors read and approved the final manuscript.

## DATA AVAILABILITY

All DNA sequencing data, assembled genomes, and annotations used for this study have been submitted to the NCBI database under SRA accession number XXXXX and BioProject accession number XXXXX.

