## Supplemental information - Figures, Texts, and Tables (captions) for "Phylogenomics provides insights into the evolution of cactophily and host plant shifts in *Drosophila*"

#### Text S1. Taxonomic classification for the 12 species included in this study.

This classification was made using formal and informal taxonomic ranges and includes only species whose genomic data were used in this project. See below the list of consulted references to generate the classification:

- https://wiki.flybase.org/wiki/FlyBase:Phylogeny
- Clark et al. (2007)
- Oliveira et al. (2012)
- Russo et al. (2013)
- Sanchez-Flores et al. (2016)
- Hurtado et al. (2019)
- Moreyra et al. (2019)
- Thurmond et al. (2019)

**Family:** Drosophilidae
**Subfamily:** Drosophilinae
**Genus:** *Drosophila*
- Subgenus *Sophophora*
+ *melanogaster* group
* *melanogaster* subgroup
***D. melanogaster***
- *Drosophila* subgenus
+ *virilis* group
* *virilis* subgroup
***D. virilis***
+ *repleta* group
* *mercatorum* subgroup
***D. mercatorum***
* *hydei* subgroup
***D. hydei***
* *mulleri* subgroup
### *mulleri* complex
@ *mulleri* cluster
***D. aldrichi***
@ *mojavensis* cluster
***D. arizonae***
***D. mojavensis***
***D. navojoa***
### *buzzatii* complex
@ *buzzatii* cluster
***D. antonietae***
***D. borborema***
***D. buzzatii***
***D. koepferae***

### Text S2. Protocols of DNA extraction for short and long sequencing technologies.

**Illumina - HiSeq 2000**
We employed the Gentra® Puregene® Cell Kit designed for *D. melanogaster* to extract genomic DNA from adults flies. DNA samples were then cleaned with phenol:chloroform and stored in liquid nitrogen at -80°C.

**PacBio RSII and Sequel I**
Note: modified AGI Drosophila Genomic DNA Extraction

- **Solutions**

- Fly-A
- Tris HCI buffer 0.1M pH 8.0
- EDTA 0.1M pH 8.0
- SDS 1%
- Qiagen ProK solution (20mg/mL)
- 4M Potassium acetate
- Chloro+isoamyl 24:1
- 2-Propanol
- 70% Ethanol
- 1X TE
- Qiagen RnaseA

- **Protocol -** 150 flies of each sex yielded ~30µg of DNA
- 50 flies per 1.5mL tube with only one sex in each tube and stored at -80°C.
- With tube on ice add 500µL Fly-A solution at room temp and 2.5µL of ProK.
- Use a blue pestle to homogenize flies by pressing flies to the bottom of the tube and twisting till most flies are not intact and the orange color appears.
- Let sit on ice till all tubes are ready.
- Incubate at 65°C for 30 minutes, gently mix halfway through.
- Cool at 37°C for 3 minutes.
- Add 2.5µL of ProK and mix.
- Incubate at 37°C for 30 minutes
- Add 70µL of 4M potassium acetate and mix.
- Incubate on ice for 30 minutes.
- Centrifuge at 4°C for 30 minutes at 18,000 rcf.
- Transfer supernatant to new 1.5mL tube be careful not to transfer debris.
- Add equal volume of chloro+isoamyl 24:1 and mix by rocking back and forth 40 times.
- Centrifuge at 4°C for 5 minutes at 10,500 rcf.
- Transfer the upper phase to a new 1.5mL tube. Careful not to get any of the interface.
- Add 350µL of 2-Propanol and slowly rock back and forth. Should see threads of precipitate at this point.
- Centrifuge at 4°C for 5 minutes at 10,500 rcf.
- Remove the liquid leaving the pellet.
- Wash pellet with 1mL of fresh room temperature 70% ETOH and rock back and forth to wash all surfaces.
- Centrifuge at 4°C for 2 minutes at 10,500 rcf.

- Remove all ETOH with a P1000 tip and then use a P10 tip to ensure full removal. Dry the pellet in the hood for 10-15 minutes.
- Resuspend with 1X TE. Have had success with using 30μL.
- Combine all tubes into a 1.5mL tube.
- Add 3μL of Qiagen RNaseA, mix gently and incubate at 37°C for 30 minutes.
- Nanodrop to check for quality.

#### Text S3. Repeat annotation protocol following the advanced repeat library construction tutorial of MAKER.

To mask genomes before gene annotation, repetitive element identification and classification were performed following the advanced repeat library construction tutorial of MAKER ver. 2.31.10 (Holt and Yandell 2011). Miniature inverted transposable elements (MITEs), as well as other small (< 2Kb) class 2 nonautonomous transposable elements (TEs), were first searched with MITE-hunter (Han and Wessler 2010) using parameters by default. Then, candidate elements with long terminal repeats (LTRs) were searched with LTRharvest (Han and Wessler 2010; Ellinghaus, Kurtz, and Willhoeft 2008) in two stages consisting of detecting recent and old divergent sequences. Accordingly, we collected recent LTR elements (≥99% similarity) with terminal repeats of 100-6000 bp long, size of the entire element ranging from 1.5 kb to 25 kb, TSD (target site duplication) of 5 bp flanking the element, and within 10 of its end. In addition, LTRdigest was applied to find elements with PPT (polypurine tract) or PBS (primer binding site) using eukaryotic tRNAs sequences obtained from the Genomic tRNA database (Chan and Lowe 2016). To ensure the correct boundary of the LTR, at least 50% of the PPT of PBS sequences had to be located in internal regions and with a maximum distance of 20 bp from the LTR. Moreover, we also performed further filtering of candidate elements to reduce the chance of detecting false positives: 1) candidate elements with more than 50 unknown bases (Ns) were removed; 2) pairwise alignments of candidate elements with their 50 bp flanking sequences were made using MUSCLE ver. 3.8.31 (Edgar 2004) and, if flank sequences were also alignable (half of the total nucleotides identical with at least 60% identity excluding gaps), the candidate element was excluded; 3) LTR sequences from candidate elements retained after the steps above were used to mask the putative internal regions of other elements and, if LTR or MITE sequences were detected in internal regions, remove those elements nested with other insertions; 4) a transposase database available at the RepeatMasker ver. 3.3.0 repository (Smit, Hubley & Green at http://www.repeatmasker.org/), was used to also search DNA transposons in internal regions and remove elements with significant matches (*e-value* = $1x10^{-10}$).

Redundancy in LTR elements was reduced by selecting only representative sequences. To this end, several rounds of BLASTN were employed in all-vs-all searches of LTR elements. The most representative element in each round was chosen using the greatest number of matches obtained considering 80% identity and 90% coverage cut-off thresholds, and retained in the final repeat library. Then, this representative sequence and its matches were excluded from the set of elements in the next round of BLASTN searches, leading to the generation of a second representative sequence. This selection process was repeated until no additional sequences could be found, and all representative elements were combined to form the set of recent LTR elements at 99% identity (LTR99). In the second stage, we repeated the LTR identification to recover old LTR retrotransposons by applying a 85% similarity cutoff (LTR85). The detected LTR85 sequences were masked with the representative LTR99 sequences and excluded if the match had 80% identity and 90% coverage.

As a way to create species-specific repeat libraries, we employed a de novo approach implemented in RepeatModeler ver. 1.0.11 (https://github.com/Dfam-consortium/RepeatModeler)

using parameters by default. The genome was masked with the MITEs, LTR99, and LTR85 libraries to detect additional repeat sequences that were classified as ‘known’ or ‘unknown’. The new unknown elements were searched against the transposase database using BLASTX and reclassified as ‘known’ if significant matches (*e-value* = $1x10^{-10}$) to a transposon superfamily were found. All repeat sequences collected at this stage were compared to a *Drosophila* protein database downloaded from FlyBase release FB2019_01 (Thurmond et al. 2019). Elements with significant hits to genes were removed with ProtExcluder ver. 1.2 (Thurmond et al. 2019; Campbell et al. 2014). After excluding the possible gene fragments, we combined MITEs, LTR99, LTR85, RepeatModeler known, and unknown repeat sequences to generate species-specific libraries.

### Text S4. Contamination removal from annotations.

To detect potential non-eukaryotic proteins, the annotated protein dataset of each species was compared to the UniProtKB release 2020_01 and eggNOG ver. 5.0 (Huerta-Cepas et al., 2019) databases. The *blastp more-sensitive* and *e-value* = $1x10^{-5}$ parameters were set in Diamond v0.9.30.131 (Buchfink et al., 2015) for mapping against UniprotKB, whereas default parameters were used for the eggNOG-mapper ver. 2 (Huerta-Cepas et al., 2017) searches in the eggNOG database. All proteins with a hit to a non-eukaryotic protein were removed from the genome annotation.

### Figures

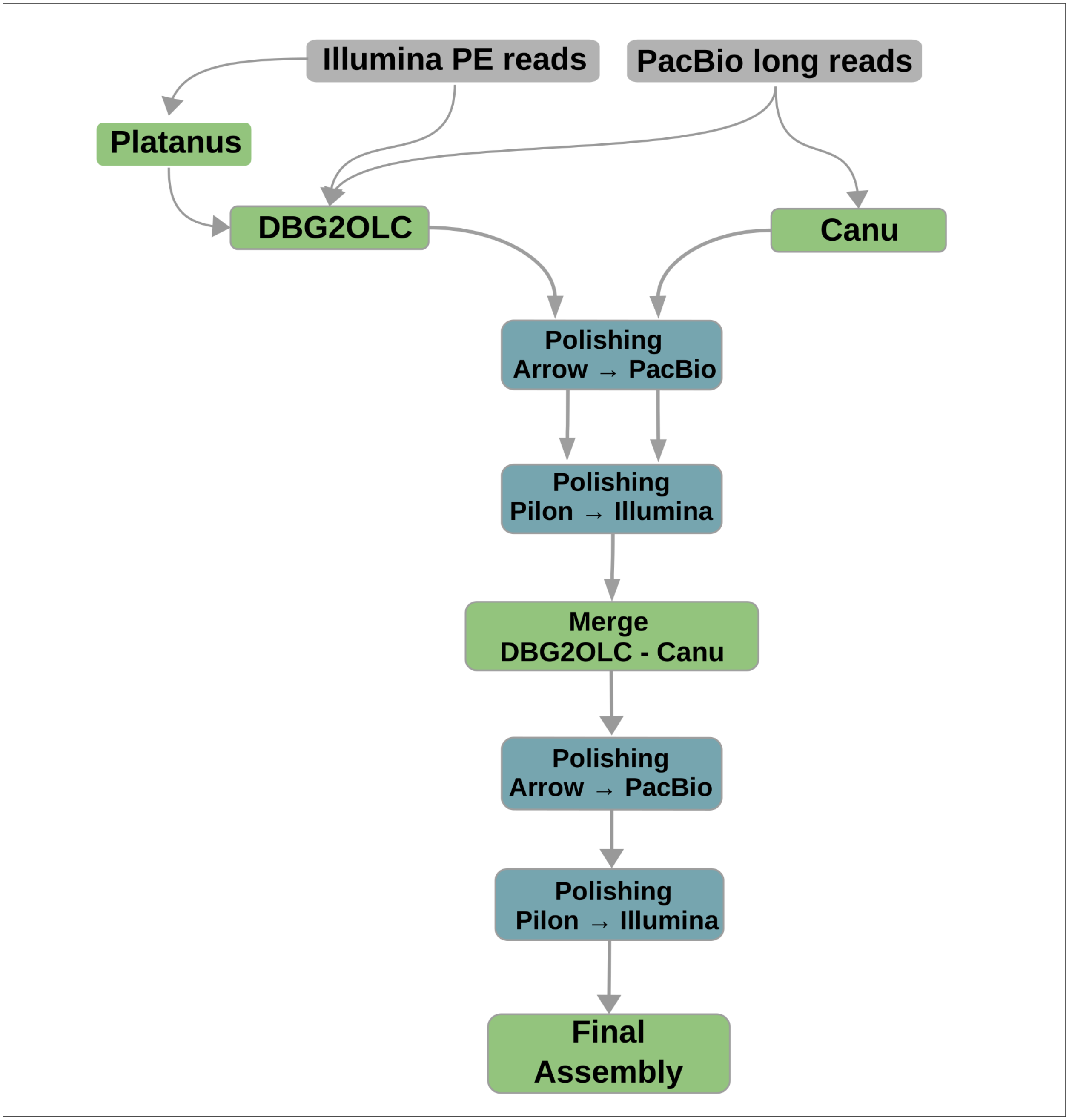


**Figure S1.** Genome assembly protocol. The pipeline involves the use of Platanus, DBG2OLC, and Canu to generate three different assemblies that are subsequently merged and polished.

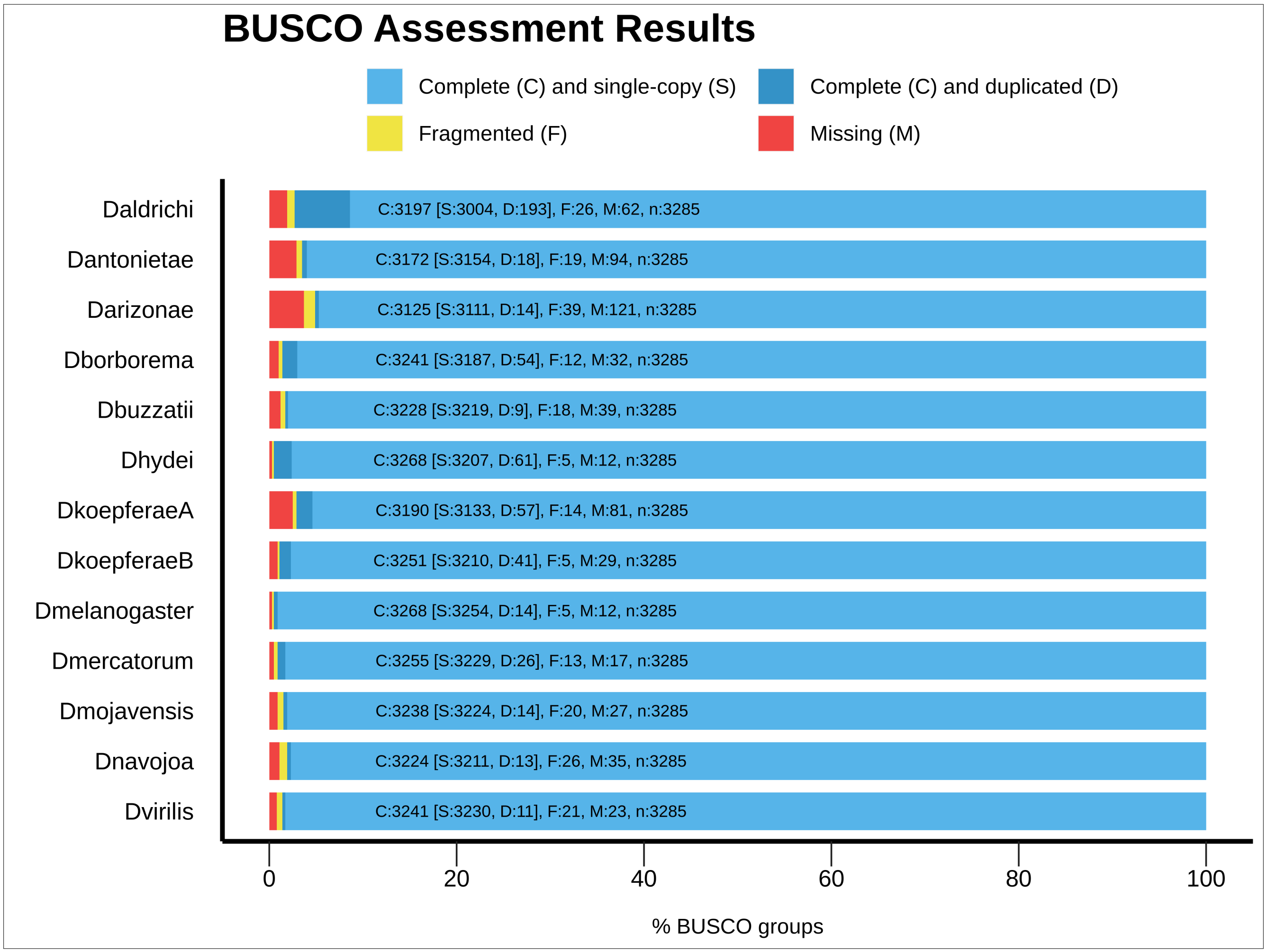


**Figure S2**. Genome completeness assessment for the thirteen Drosophila genome assemblies using 3285 dipteran BUSCO groups.

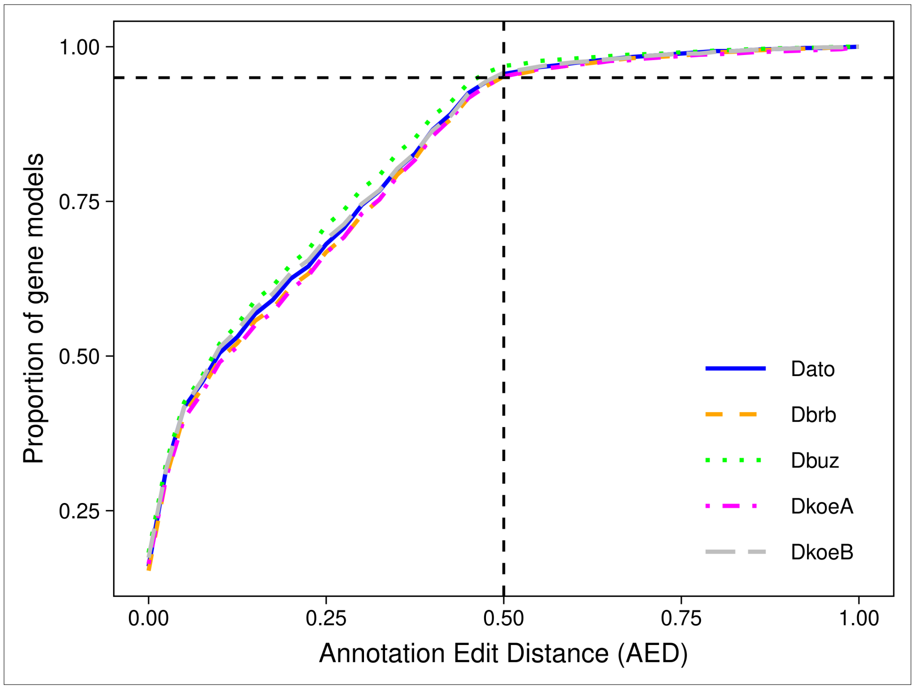


**Figure S3**. AED score distribution for genome annotations of species of the buzzatii cluster.

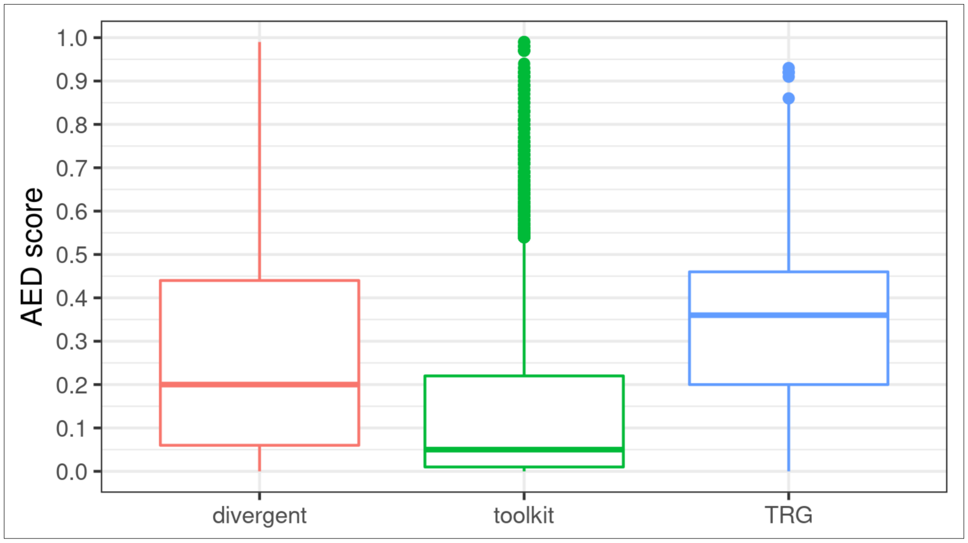


**Figure S4**. AED score distribution for candidate TRGs (divergent and validated) and toolkit genes.

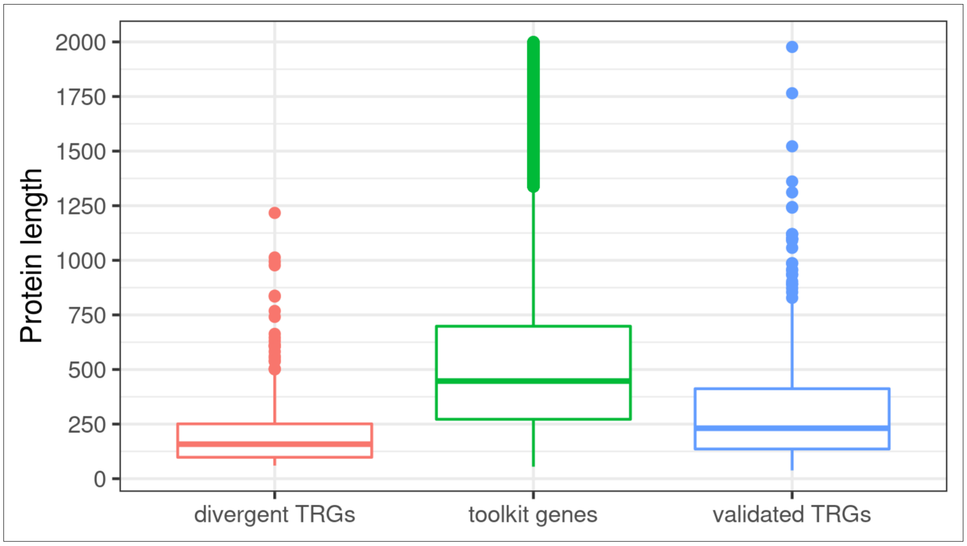


**Figure S5**. Protein length distribution for candidate TRGs (divergent and validated) and toolkit genes. Proteins of length greater than 2000 were excluded from the illustration to improve the visualization of differences between sets.

### Tables

**All tables are attached in one Microsoft Excel file.**

**Table S1**. Genome assembly information. The citation (DOI), the accession number (with version), database, and direct link are shown for each species genome.

**Table S2**. Sequencing yields and statistics for each genome.

**Table S3.** Scaffold vs contig contiguity. Score for completeness are also shown.

**Table S4**. Genome repeat content of species of the D. serido sibling set. Repeat content was assessed using the species-specific libraries and the reference Drosophila repeat library available at Dfam.

**Table S5**. Genome annotation summary. Numbers of genes, transcripts, and proteins as well as the average gene length are given for each genome.

**Table S6**. Gene- and site-concordance factors for each branch in the species tree. Values are also shown for discordance factors.

**Table S7**. Mean age estimated for each node of the species tree using two approaches.

**Table S8**. Predicted orthogroups (OGs) for the 13 Drosophila species included in this study. In each row are listed the orthogroup name and the homolog proteins that comprise it.

**Table S9**. Classification of candidate TRGs after validation. Results for the validation of incomplete TRGs within the buzzatii cluster are also shown.

**Table S10**. Molecular evolution analysis for candidate TRGs. For each candidate TRG is shown the likelihood of each model (M7, M8, and M8a), the p-values of model comparisons, and the result of the Bonferroni correction for multiple comparisons.

**Table S11**. Functional prediction of candidate TRGs. Results of the enrichment analyses performed with GOseq are shown for all annotated GOs in each branch of the species tree. In each row, a GO is shown together with the candidate TRG in which it was annotated.

**Table S12**. Functional results after reduction of redundancy in GO terms. In each row, a GO is detailed together with the candidate validated and/or divergent TRGs in which it was annotated.
